# PPARδ restrains the suppression function of intra-tumoral Tregs by limiting CIITA-MHC II expression

**DOI:** 10.1101/2024.12.16.628819

**Authors:** Qiyuan Yang, Yuqiong Liang, Tomoko Inoue-Hatanaka, Zhiqian Koh, Nadja Ilkenhans, Ethan Suman, Jingting Yu, Ye Zheng

**Affiliations:** NOMIS Center for Immunobiology and Microbial Pathogenesis, Salk Institute for Biological Studies, La Jolla, CA, USA; Razavi Newman Integrative Genomics and Bioinformatics Core, Salk Institute for Biological Studies, La Jolla, CA, USA

## Abstract

Regulatory T cells (T_reg_ cells) play a critical role in suppressing anti-tumor immunity, often resulting in unfavorable clinical outcomes across numerous cancers. However, systemic T_reg_ depletion, while augmenting anti-tumor responses, also triggers detrimental autoimmune disorders. Thus, dissecting the mechanisms by which T_reg_ cells navigate and exert their functions within the tumor microenvironment (TME) is pivotal for devising innovative T_reg_-centric cancer therapies. Our study highlights the role of peroxisome proliferator-activated receptor β/δ (PPARδ), a nuclear hormone receptor involved in fatty acid metabolism. Remarkably, PPARδ ablation in T_reg_ escalated tumor growth and augmented the immunosuppressive characteristics of the TME. This absence of PPARδ spurred an increased expression of genes central to antigen presentation, notably CIITA and MHC II. Our results showcase a novel association where the absence of CIITA in PPARδ-deficient T_reg_ bolsters anti-tumor responses, casting CIITA as a pivotal downstream regulator of PPARδ within T_reg_. *In vitro* assays demonstrated that elevated CIITA levels enhance the suppressive capacity of T_reg_, facilitated by an antigen-independent interaction between T_reg_-MHC II and T_conv_-TCR/CD4/Lag3. A significant revelation was the role of type 1 interferon as a TME signal that promotes the genesis of MHC II^+^ T_reg_; PPARδ deficiency intensifies this phenomenon by amplifying type 1 interferon signaling, mediated by a notable upsurge in JAK3 transcription and an increase of pSTAT1-Y701. In conclusion, the co-regulation between TME cues and PPARδ signaling shapes the adaptive and suppressive roles of T_reg_ cells through the CIITA-MHC II pathway. Strategically targeting the potent MHC II^+^ T_reg_ population could open a new avenue for cancer therapies by boosting anti-tumor defenses while curbing autoimmune threats.

**Highlights:**

- PPARδ T_reg_ conditional knockout mice show accelerated tumor growth due to increased expression of CIITA-MHC II.
- Type I interferon signal regulates T_reg_ CIITA-MHC II axis *in vitro* and *in vivo*.
- PPARδ attenuates Type I interferon response and restrains CIITA-MHC II expression in T_reg_ cells.
- T_reg_ suppressive function is enhanced by T_reg_ MHC II’s direct interaction with TCR/CD4/Lag3 on T_eff_ cells.

## Introduction

Regulatory T cells (T_reg_ cells) are essential in controlling hyperactive immune responses and significantly influence cancer progression in patients(*1–5*). Multiple studies employing various methods to disable or deplete T_reg_ cells have demonstrated that they could be a potential target to enhance anti-tumor immunity.(*6–8*) However, systemic removal of T_reg_ cells not only boosts anti-tumor responses but also increases the risk of autoimmunity (*1, 8–12*). This dual effect underscores the need to unravel the mechanisms by which T_reg_ cells navigate and exert their functions within the tumor microenvironment (TME) in order to develop safer and more effective T_reg_-centric cancer therapies.

Extensive research has illustrated how T_reg_ cells maintain tissue homeostasis and suppress the function of effector T cells (T_eff_ cells) in both physiological and pathological conditions(*13–16*). Current models suggest that T_reg_-mediated suppression operates through both cell contact-dependent(*17–22*) and cell contact-independent mechanisms(*23–28*). Despite these advances, key questions remain regarding how T_reg_ adapts to local environments, especially the tumor microenvironment (TME), as well as how T_reg_ cells suppress intra-tumoral T_eff_ cells. The mechanism by which T_reg_ cells shape and contribute to tumor progression needs further elucidation.

Accumulated evidence indicates that tumor-infiltrated T cells undergo metabolic rewiring, particularly in lipid metabolism, to adapt to the TME(*29–32*). These metabolic adaptations are regulated by a group of transcription factors, including the nuclear receptor family known as Peroxisome Proliferator-activated receptors (PPARs)(*33*). Among the three PPAR isoforms, PPARδ is ubiquitously expressed and plays a critical role in lipid metabolism(*34*), inflammation(*35*), cellular survival(*36*), differentiation(*37, 38*), as well as maintaining energy balance in various tissues(*39*). In T cells, PPARδ activation has been reported to restrict Th1 and Th17 responses while promoting Th2 responses(*40–42*). However, the role of PPARδ in T_reg_ cells was not clearly established.

In this study, we employ conditional knockout PPARδ mice to delve into the function of PPARδ in T_reg_ cells in anti-tumor immunity. PPARδ ablation in T_reg_ enhances tumor growth and increases the immunosuppressive nature of the TME by activating the CIITA-MHC II axis in intra-tumoral T_reg_. Type 1 interferon signaling emerges as a key driver of MHC II upregulation in intra-tumoral T_reg_ cells, while PPARδ restrains MHC II expression through weakening the JAK-STAT pathway.

## Results

### Ablation of PPARδ in T_reg_ cells leads to accelerated tumor growth

Among the PPAR family members, PPARδ is the only one consistently expressed throughout all stages of mouse T-cell differentiation (**Fig. S1a**). We assessed the expression level of PPARα, PPARγ, and PPARδ in peripheral T cell subsets including CD8 T cells, conventional T cells (T_conv_), and T_reg_ cells isolated from the peripheral lymph nodes. PPARδ messenger RNA levels were the highest among the PPAR isoforms (**Fig. S1b-d**), suggesting it may play a significant role in regulating T cell function. To investigate PPARδ’s potential in modulating T_reg_ cell function, we generated T_reg_-specific PPARδ conditional knockout (PPARδ cKO) mice by crossing the *Foxp3*^YFP-cre^ mouse with the *PPARδ*^fl/fl^ mouse(*43*). These PPARδ cKO mice had normal distributions of CD4^+^ T cell, CD8^+^ T cell, and T_reg_ cell populations in the thymus and the periphery with unaffected Foxp3 expression levels in T_reg_ cells (**Fig. S2a-f**). Specific ablation of PPARδ in T_reg_ cells did not induce spontaneous activation in conventional T cells (**Fig. S2g,h**). Further, The activation status of peripheral CD4^+^ and CD8^+^ T cells in PPARδ cKO mice is comparable to WT controls based on IFNγ, IL-4, IL-13, and IL-17A expression (**Fig. S2i-n**), implying that Treg expression of PPARδ is dispensable for immune homeostasis at steady state. We performed Treg cell apoptosis and proliferation assays to further characterize PPARδ’s role in Tregs and observed comparable results between wild-type (WT) T_reg_ cells and PPARδ-deficient T_reg_ cells (**Fig. S3a-f**). To evaluate whether PPARδ deficiency affects Treg’s metabolic function, we performed fatty acid or glucose uptake assays and measured the mitochondrial membrane potential of cultured Tregs, and observed similar results between WT and PPARδ-deficiency T_reg_ cells (**Fig. S4a-e**). In conclusion, endogenous PPARδ seems to play a minimal role in regulating T_reg_ cell homeostasis and function, including maintaining mitochondrial function, lipid and glucose metabolism, proliferation, apoptosis, differentiation, and T_reg_ cells’ suppressive function in the steady states.

A large body of studies showed that T_reg_ cells can be enriched in tumors and suppress anti-tumor immunity (*44–46*). We tested whether PPARδ regulates T_reg_ cell adaptation and function in cancer immunity by inoculating tumor cells in WT and PPARd cKO mice and tracking tumor growth. Remarkably, PPARδ cKO mice exhibited an increase and acceleration in the growth of tumors compared to their WT counterparts after implanting the B16 melanoma, the MC38 adenocarcinoma, and the EL4 thymoma cells (**Fig. 1a**). Immunoprofiling across the three tumor types revealed modest changes in the percentages of intra-tumoral innate immune cell subsets (**Fig. S5**), CD4^+^ conventional T cells, and CD8^+^ T cells (**Fig. 1b**). The percentages of T_reg_ among tumor-infiltrated CD4^+^ T cells were consistently higher in PPARd cKO mice compared to WT mice (**Fig. 1c**). Notably, PPARδ cKO mice displayed a more immune suppressive TME, as evidenced by a decrease in IFNγ^+^ and TNFα^+^ tumoricidal subsets among tumor-infiltrating CD4^+^ conventional T cells and CD8^+^ T cells (**Fig. 1d,e**). Thus, PPARδ restricts intra-tumoral T_reg_ number and function, leading to enhanced anti-tumor immunity.

**Figure 1:**
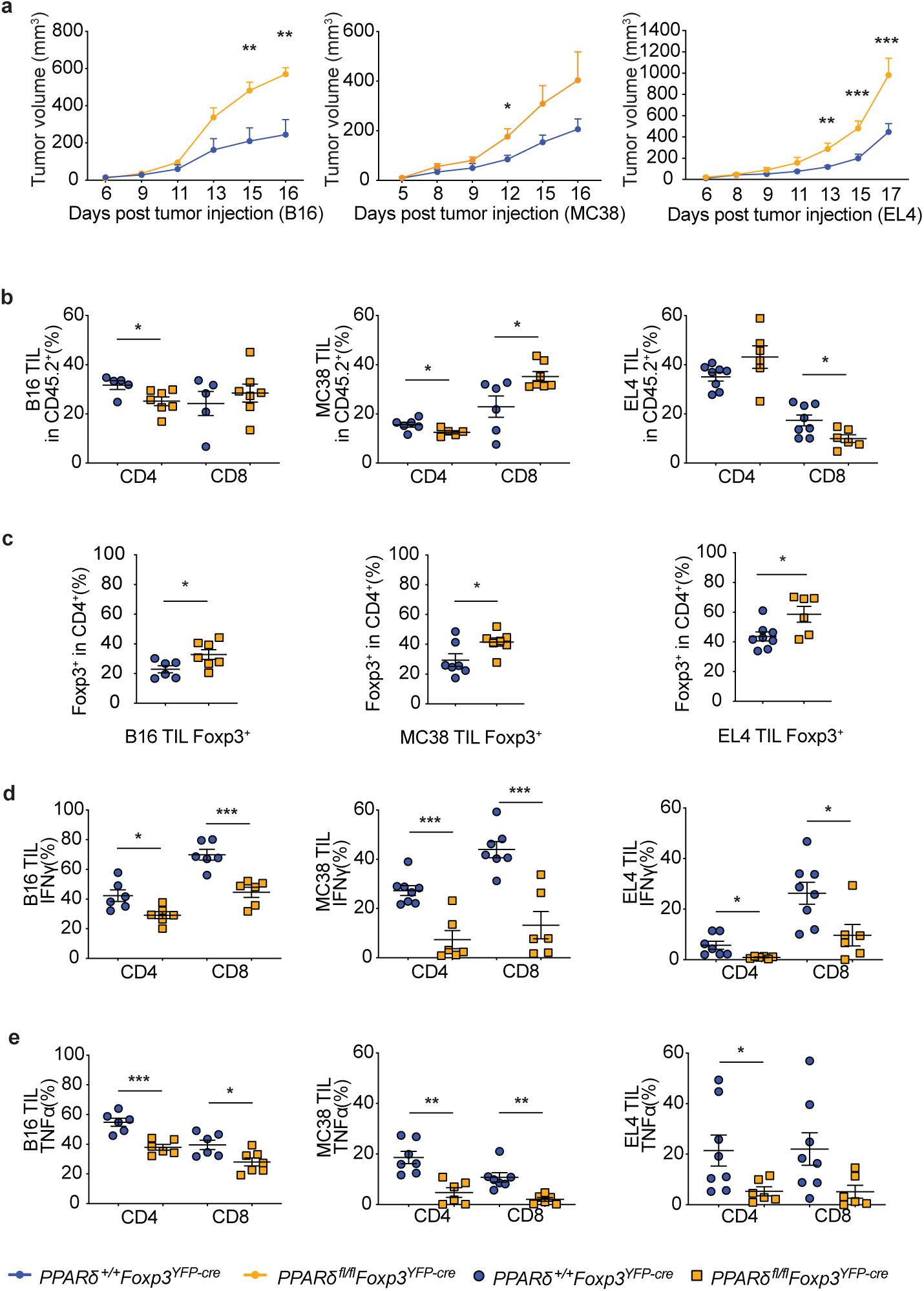
Regulatory roles of PPARδ in T_reg_ cells modulate anti-tumor immunity and tumor growth dynamics. A | Growth trajectories of B16F10 melanoma, MC38 colon cancer, and EL4 thymoma in PPARδ-sufficient (*PPARδ^+/+^Foxp3^cre^*) and PPARδ-deficient (*PPARδ^fl/fl^Foxp3^cre^*) hosts, measured by tumor volume over time post-inoculation. For each tumor model, 5 x 10^5^ tumor cells were inoculated subcutaneously. B | Differential composition of CD4^+^ and CD8^+^ T cells within the TIL population across B16, MC38, and EL4 tumor models in both genotypes. C | Proportion of Foxp3^+^ cells within the CD4^+^ TIL compartment, comparing T_reg_ prevalence in the tumor milieu between the two genotypes. D | IFNγ expression profiles in CD4^+^ and CD8^+^ TILs, indicating cytokine-mediated immune responsiveness. E | TNFα expression levels in CD4^+^ and CD8^+^ TIL subsets, reflecting pro-inflammatory response modulation. *The data represent a synthesis of 2-4 independent experiments for each tumor type, involving 6-8 mice per group. Statistical analyses were performed using two-tailed unpaired t-tests, with significance denoted as: *P < 0.05, **P < 0.01, ***P < 0.001. Data are expressed as mean values ± SEM. MFI stands for mean fluorescence intensity*.

### The CIITA-MHC II axis is regulated by PPARδ in T_reg_ cells

To elucidate how PPARδ in intra-tumoral T_reg_ restricts tumor growth, we performed RNA-sequencing (RNA-seq) experiments on intra-tumoral T_reg_ cells isolated from PPARδ cKO and WT control mice. RNA-seq data analysis uncovered 92 and 56 differentially expressed genes (DEGs) in the B16 and MC38 tumor models, respectively (**Fig. 2a,b**). Notably, among these DEGs, a group of the MHC II related genes were significantly upregulated in PPARδ-deficient intra-tumoral T_reg_ cells (**Fig. 2a,b**). In an analysis of the upregulated DEGs from both B16F10 and MC38 tumor models, there are 10 overlapping genes, half of which are MHC II related genes (**Fig. 2c**). Consistently, over-representative analysis (ORA) of the DEGs revealed enrichment of genes involved in the antigen processing and presentation pathway in PPARδ-deficient T_reg_ cells from both B16F10 and MC38 tumors (**Fig. 2d**). While mouse T cells typically do not express MHC II, which is highly expressed by antigen-presenting cells such as dendritic cells, macrophages, as well as certain tissue-specific with functional roles, such as mast cells, basophils, eosinophils, ILC3s, and microglia(*47*). We observed an increase in the percentage of MHC II^+^ T_reg_ cells and the mean fluorescent intensity (MFI) representing MHC II expression level in T_reg_ cells in B16, MC38, and EL4 tumors (**Fig. 2e**). In addition to MHC II genes, we also found a group of antigen presentation related genes upregulated in the PPARδ-deficient intra-tumoral T_reg_ cells, including their master regulator CIITA(*48*), H2-O, H2-DM, and Cd74 (**Fig. 2f,g**). This implies MHC II^+^ T_reg_ cells could be equipped with the antigen presentation machinery. Of note, although the expression level of MHC II in T_reg_ is much higher than CD4 conventional cells, CD8 T cells, and NK cells, it is substantially lower than professional APCs such as dendritic cells, macrophages, and B cells (**Fig. S6**), hinting that the MHC II expressed in these T_reg_ cells might carry a function that is different from classic antigen presentation cells.

**Figure 2:**
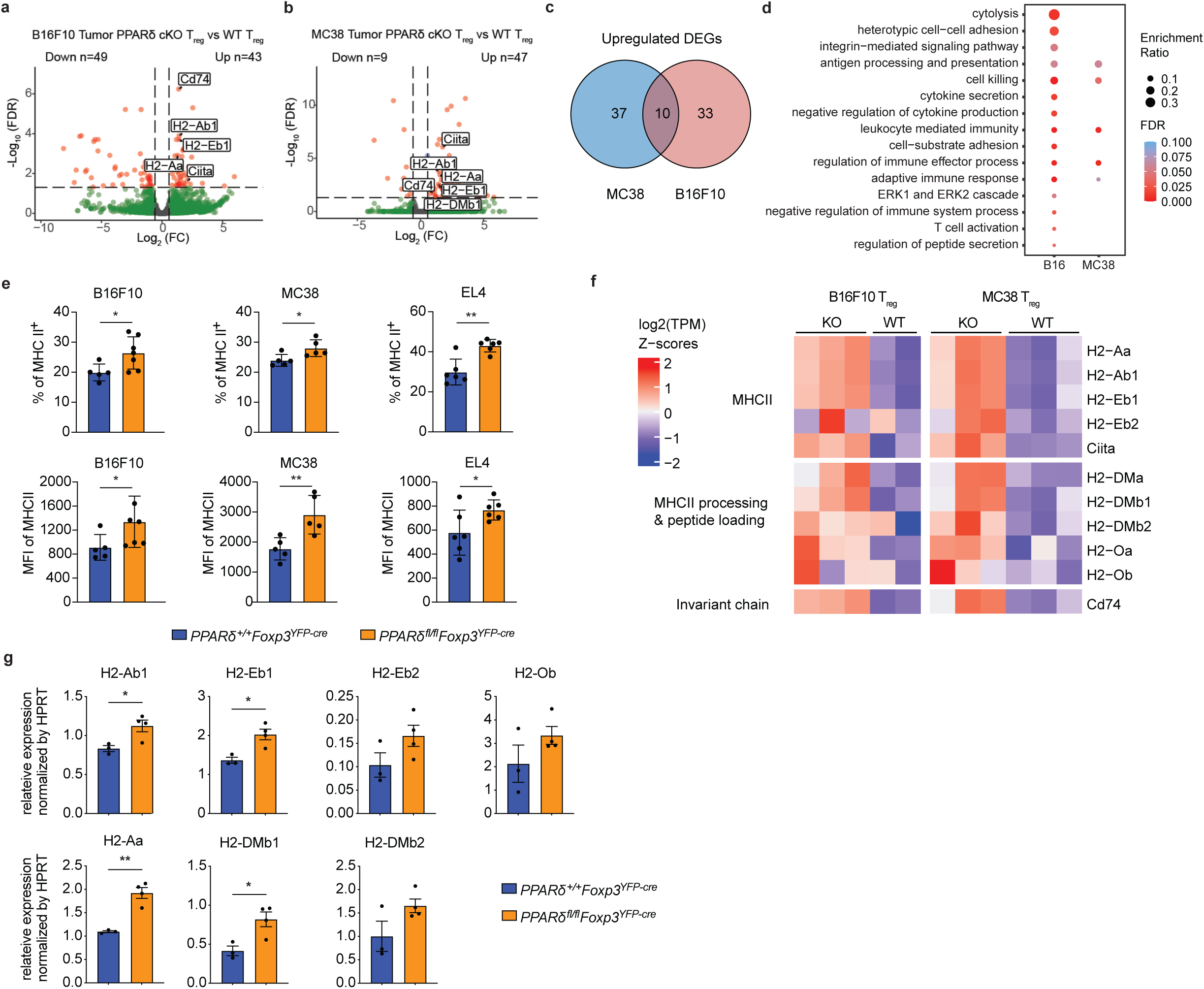
Enhanced expression of MHC II genes in PPARδ-deficient intratumoral T_reg_ cells. A and B | Scatter plots displaying differentially expressed genes in T_reg_ cells extracted from B16F10 melanoma (A) and MC38 colon carcinoma (B) in PPARδ-deficient mice compared to controls. Genes meeting the FDR<0.05 and fold change>1.5 criteria are shown, with a particular emphasis on MHC II-related genes. C | Venn diagram demonstrating the overlap of upregulated differentially expressed genes (DEGs) between B16F10 and MC38 T_reg_ cells in PPARδ-deficient mice. D | Over-Representation Analysis (ORA) for upregulated genes in T_reg_ cells from B16F10 tumors, applying an FDR<0.05 (dark blue) and fold change>1.5, conducted using WebGestalt. E | Flow cytometric quantification of MHC II expression in intratumoral T_reg_ cells from *PPARδ^+/+^Foxp3^cre^* and *PPARδ^fl/fl^Foxp3^cre^*mice, indicating upregulation in the absence of PPARδ. F | Heatmap representing expression profiles of genes associated with the MHC II antigen presentation pathway in T_reg_ cells from both B16F10 and MC38 tumors, comparing *PPARδ^+/+^Foxp3^cre^*and *PPARδ^fl/fl^Foxp3^cre^* genotypes. G | Quantitative RT-PCR analysis of MHC II gene expression in sorted, purified T_reg_ cells from B16F10 tumor-bearing C57BL/6 mice, including *PPARδ^+/+^Foxp3^cre^*and *PPARδ^fl/fl^Foxp3^cre^* mice, aged 8-12 weeks. Relative expression levels of certain genes are normalized by HPRT. *All analyses were based on gene expression data that passed a threshold of FDR<0.05 and fold change>1.5. The heatmap z-scores represent expression levels normalized across all samples. Data are presented as mean values ± SEM. Statistical analysis was performed using an unpaired, two-tailed t-test, with significance indicated as: *P < 0.05, **P < 0.01. ns denotes not significant. MFI stands for mean fluorescence intensity*.

To test whether the up-regulation of CIITA/MHC II in PPARδ cKO T_reg_ cells contributes to the suppression of tumor immunity, we performed a T cell adoptive transfer experiment. Rag1 KO mice were transferred with Ly5.1 CD4 and CD8 naïve T cells along with T_reg_ cells transduced with sgRNA targeting PPARδ, sgRNAs targeting both PPARδ and CIITA, or a non-targeting control sgRNA. These mice were inoculated with B16F10 melanoma cells on the next day and monitored for tumor growth (**Fig. 3a**). The double knockout group showed a significant reduction of tumor growth compared to the PPARδ single knockout group, and it is comparable to the control group (**Fig. 3b**). Flow cytometry analysis confirmed the reduction of MHC II expression in sgCIITA transduced T_reg_ cells (**Fig. 3c**). Next, we analyzed cytokine expression in the intra-tumoral T cells. Knockdown of CIITA and PPARδ in T_reg_ cells restored IFNγ and TNFα expression in both CD4 and CD8 T cells compared to mice with PPARδ single knockdown (**Fig. 3d-g**). Therefore, the up-regulation of the CIITA/MHC II is a major contributor to the increase in immune suppression in PPARδ cKO mice.

**Figure 3:**
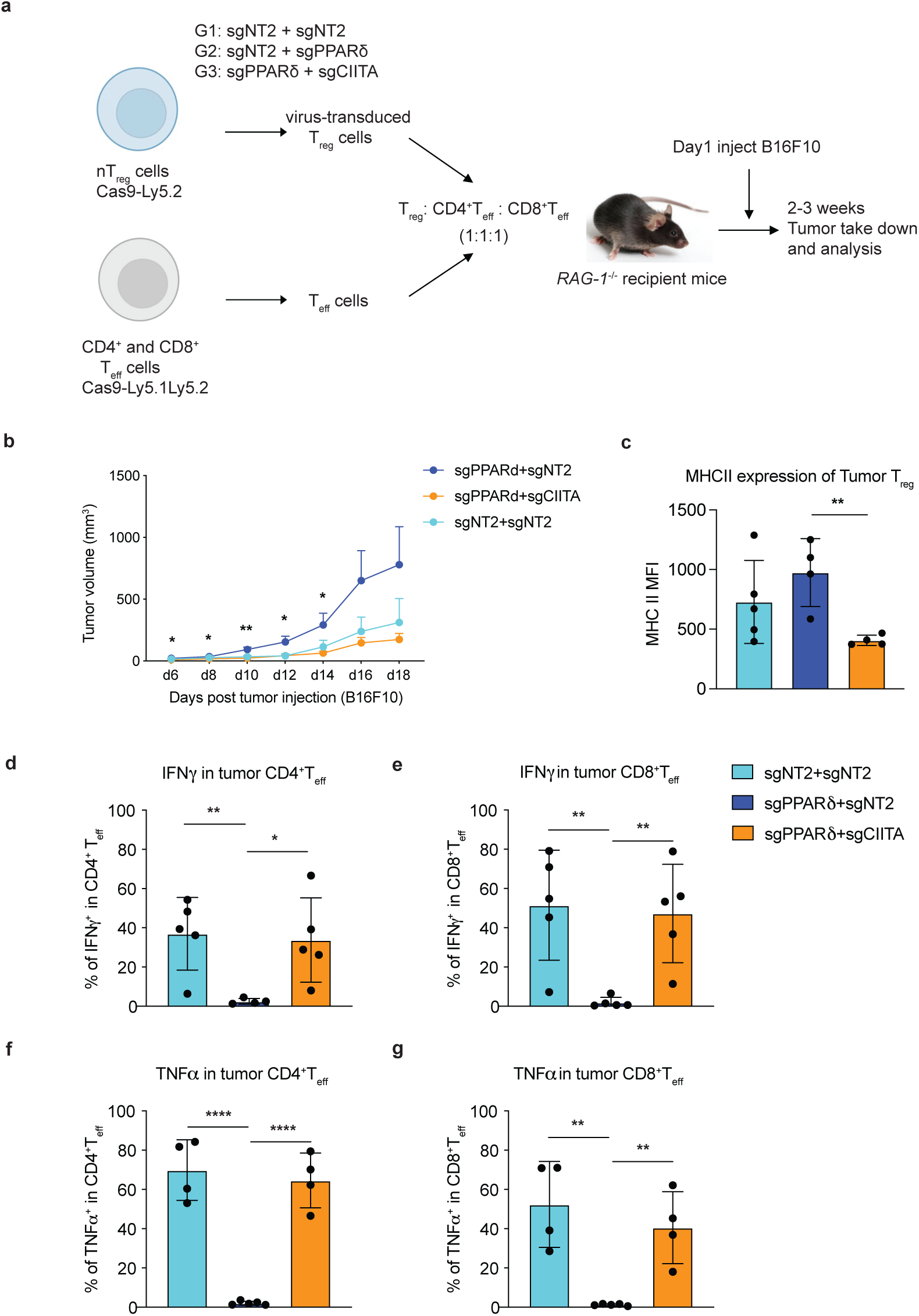
CIITA-MHC II axis is downstream of PPARδ signaling in intratumoral T_reg_ cells. A | Schematic diagram of adoptive T cell transfer tumor model. B | Tumor growth curve of T cell adoptive transfer *Rag1*^-/-^ recipient mice receiving 2.5 × 10^5^ B16F10 melanoma cells. Statistical analysis was shown between PPARδ single knockout group and PPARδ & CIITA double knockout group. C | Flow cytometric analysis of MHC II expression level in transferred T_reg_ cells from tumors. D-G | IFNγ (D, E) and TNFα (F, G) cytokine staining of intra-tumoral CD4 and CD8 T_eff_ cells. *Statistical analysis was performed using an unpaired, two-tailed t-test, with significance indicated as: *P < 0.05, **P < 0.01, ****P < 0.0001. MFI stands for mean fluorescence intensity*.

### Type I interferon signal regulates intra-tumoral T_reg_ CIITA-MHC II axis

To delineate how PPARδ suppresses CIITA-MHC II, we need to first examine how these genes are up-regulated in intra-tumoral T_reg_ cells. To this end, we treated T_reg_ cells with a group of pro-inflammatory cytokines, including IL-6, IFN-α, IFN-β, IFN-γ, and TNFα, and measured MHC II expression (**Fig. S7a, b**). Only type 1 interferons (IFN-α and IFN-β) up-regulated MHC II expression in T_reg_ cells. In PPARδ deficient T_reg_ cells, type 1 interferons induced MHC II expression at higher levels compared to WT T_reg_ (**Fig. 4a, b**). To test whether blocking the type 1 interferon pathway would reduce the intra-tumoral MHC II^+^ T_reg_ cell population, we conducted an adoptive T cell transfer tumor growth experiment, comparing T_reg_ cells knockout of type 1 interferon receptor IFNAR1 to WT control T_reg_ cells (**Fig. S7c**). Although the difference in tumor volume between the IFNAR1 knockout group and the control group was not significant (**Fig. S7d**), we observed a marked reduction in MHC II expression level in IFNAR1 knockout T_reg_ cells compared to control T_reg_ cells. Interestingly, this difference was only observed in tumor-infiltrated T_reg_ cells, not in the spleen, suggesting that type 1 interferon is a primary signal in the tumor microenvironment to induce MHC II expression in T_reg_ cells (**Fig. S7e-h**).

**Figure 4:**
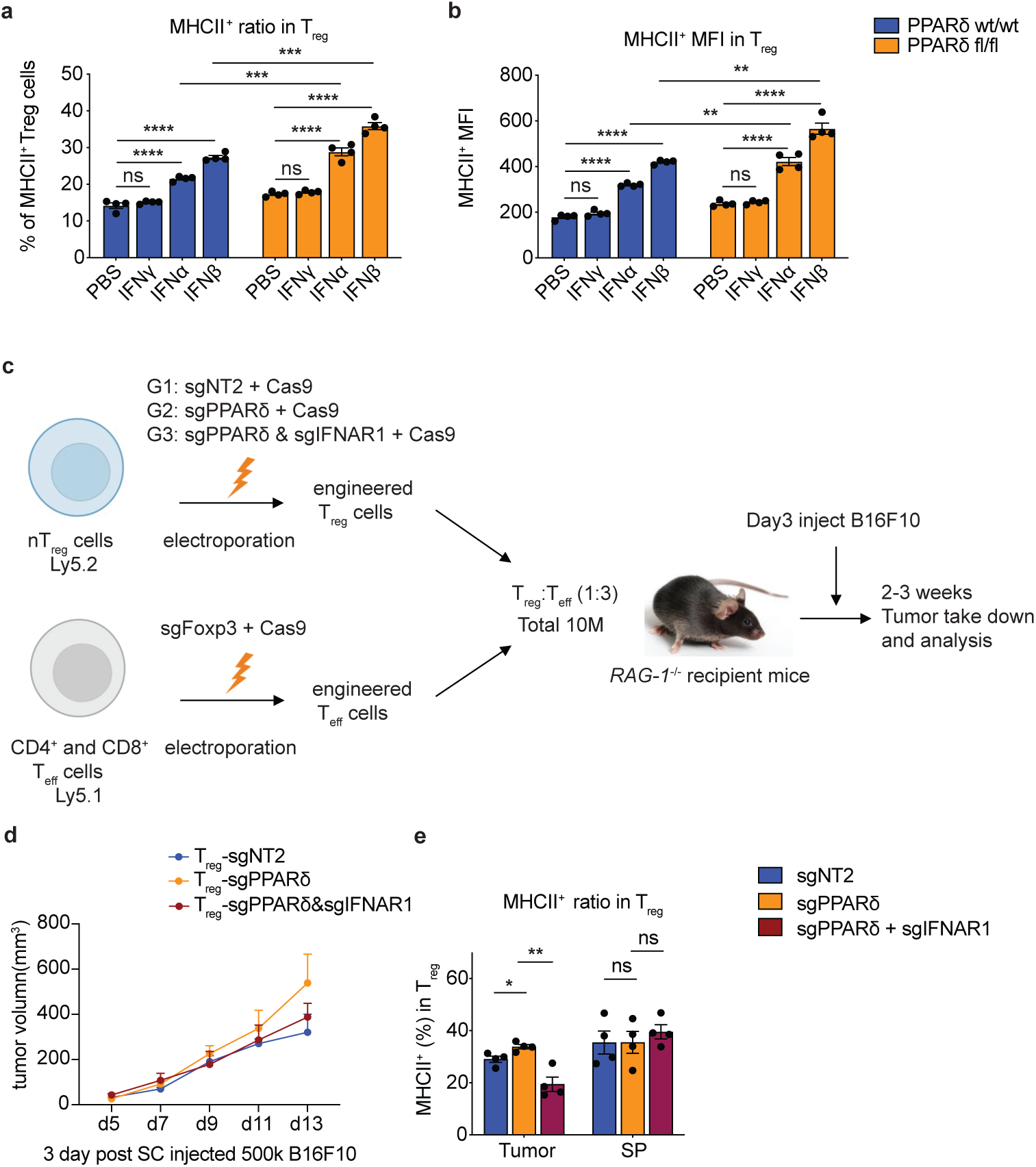
Type I interferon signaling regulates intratumoral T_reg_ CIITA-MHC II axis. A | Flow cytometric analysis of MHC II expression on T_reg_ cells from PPARδ wild-type and conditional knockout mice after cytokine stimulation with IFN-γ, IFN-α, and IFN-β compared to PBS control. B | Graphical representation of the mean fluorescence intensity (MFI) of MHC II on T_reg_ cells following the same treatments as in (A). C | Schematic overview of the experimental setup for adoptive transfer of electroporated T_reg_ cells and effector T cells (T_eff_) into B16F10 melanoma-bearing Rag1-/- mice. The diagram details the groups: sgNT2+Cas9 (control), sgPPARδ+Cas9, and sgPPARδ & sgIFNAR1+Cas9. D | Tumor growth curves for B16F10 melanoma in Rag1-/- recipient mice that received engineered T_reg_ cells according to the schematic in (C), measured over time post subcutaneous tumor cell injection. E | Flow cytometric quantification of the percentage of MHC II^+^ T_reg_ cells within the tumor and spleen (SP), comparing the outcomes among various genetically engineered T_reg_ groups. Data are presented as mean values ± SEM. Statistical analysis was performed using an unpaired, two-tailed t-test, with significance indicated as: *P < 0.05, **P < 0.01, ***P < 0.001, ****P < 0.0001, ns denotes not significant. MFI stands for mean fluorescence intensity.

To further investigate the relation between type 1 interferon signaling and PPARδ pathway, and their influence on the CIITA-MHC II axis, we performed an adoptive T cell transfer tumor experiment using T_reg_ cells knockdown of both IFNAR1 and PPARδ (**Fig. 4c**). Knockdown of IFNAR1 downregulated MHC II expression to baseline levels in PPARδ-deficient T_reg_ cells (**Fig. 4d,e**), suggesting type 1 IFNs are the primary signals driving MHC II expression in T_reg_ cells, with PPARδ and type 1 IFN convergently regulate CIITA-MHC II axis.

### PPARδ suppresses MHC II expression through the Jak3/Stat1 signaling pathway

To elucidate how PPARδ modulates the CIITA-MHC II axis, we performed PPARδ Cut-and-Run experiment in T_reg_ cells expressing a TY1-tagged PPARδ with or without the PPARδ agonist GW501516 (**Fig. S8a**). We observed a successful agonist treatment, demonstrated by a substantial overlap of peaks and a significant increase in the number of PPARδ binding peaks (**Fig. S8b**). This assay unveiled binding peaks at the established PPARδ target genes, such as Plin2, Pdk4, Angptl4, and Cpt1a, underscoring PPARδ’s influence on lipid metabolism-related genes (**Fig. S8c-f**). However, the absence of PPARδ binding to class II genes and CIITA loci suggests that PPARδ regulates these genes in an indirect manner (**Fig. S8g,h**). By comparing PPARδ-regulated genes from the Cut-and-Run assay and the DEGs from RNA-seq of the cKO and WT tumor-infiltrating T_reg_ cells, we identified JAK3 as a potential PPARδ direct target with enhanced expression in cKO Tregs compared to WT T_reg_ cells (**Fig. 5a-c**). Based on this result, we hypothesized that PPARδ may influence class II gene expression by transcriptionally repressing JAK3 expression, thereby weakening Stat1 phosphorylation and type 1 interferon signaling. To substantiate our hypothesis, freshly isolated splenocytes and lymphocytes were treated briefly under various conditions—with or without IFN-β, with or without the JAK3-specific inhibitor WHI-P131—and assessed pSTAT1-Y701 phosphorylation levels. Our analyses revealed that JAK3 inhibition corresponds with a decreased pSTAT1-Y701 level (**Fig. 5d,e**), suggesting that in T_reg_ cells, type 1 IFN activates Stat1 through JAK3. Stat1 phosphorylation levels in PPARδ cKO T_reg_ cells is higher than WT T_reg_ cells, suggesting that PPARδ suppresses class II gene expression by inhibiting Jak3 expression.

**Fig. 5.**
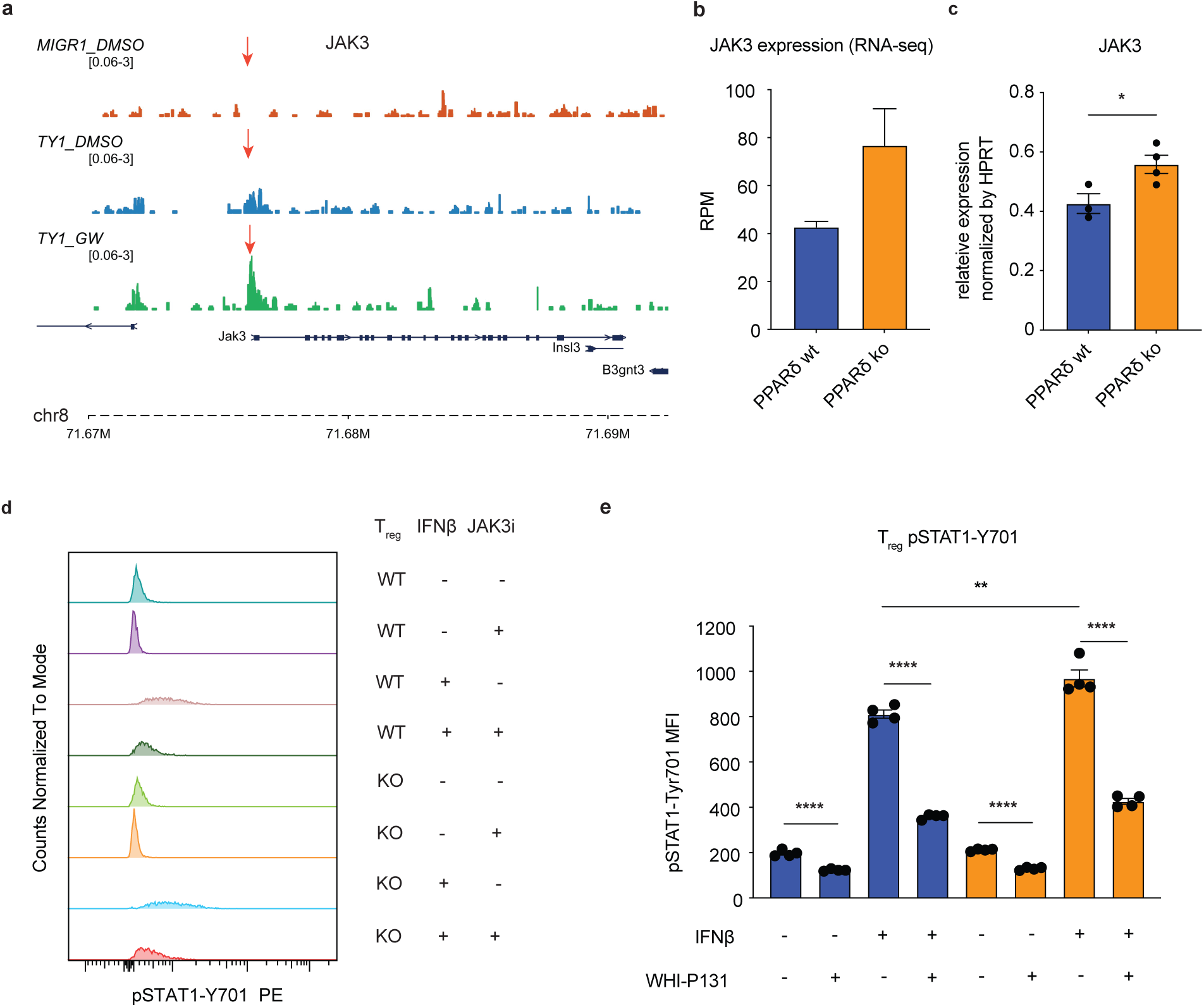
PPARδ as a Transcriptional Repressor of Jak3, Inhibiting Stat1 Phosphorylation. A | Cut-and-Run sequencing tracks showing the binding of PPARδ at the JAK3 locus in T_reg_ cells, comparing PPARδ wild-type and knockout cells (with or without GW501516). Red arrows point to the PPARδ binding locus. B | RNA-seq analysis depicting JAK3 expression levels in T_reg_ cells from tumor tissue, highlighting differences between PPARδ wild-type and knockout cells. C | Quantitative RT-PCR analysis of JAK3 expression in sorted, purified T_reg_ cells from naive C57BL/6 mice both *PPARδ^+/+^Foxp3^cre^* and *PPARδ^fl/fl^Foxp3^cre^ genotypes*, aged 8-12 weeks. The relative expression level is normalized by HPRT. D | Histograms representing intracellular staining for phosphorylated STAT1 (pSTAT1-Y701) in T_reg_ cells derived from splenocytes of *PPARδ^+/+^Foxp3^cre^*and *PPARδ^fl/fl^Foxp3^cre^* mice treated with IFN-β or JAK3 inhibitor WHI-P131. E | Statistical analysis correlating to (D), showing mean fluorescence intensity (MFI) data for pSTAT1-Y701 in the different treatment groups. *Statistical analysis was performed using an unpaired, two-tailed t-test, with significance indicated as: *P < 0.05, **P < 0.01, ****P < 0.0001. Normalized counts are shown for pSTAT1-Y701 phosphorylation under the various treatment conditions.* T_reg_ cells *were assessed for response to IFN-β stimulation or inhibition via JAK3-specific inhibitor, WHI-P131*.

### Increased CIITA/MHCII expression in T_reg_ cells enhances T_reg_ suppressive function *in vitro*

Based on our observations of accelerated tumor growth, we hypothesized that MHC II expression enhances T_reg_ cell’s suppressive function, and a more suppressive TME correlates with a higher MHC II^+^ T_reg_ proportion in total T_reg_ cells. To validate this, we performed an *in vitro* suppression assay (IVSA), measuring the suppressive function of MHC II^high^ and MHC II^low^ T_reg_ cells. To test whether MHC II^+^ T_reg_ cells directly interact with T_eff_ cells and act as APCs (**Fig. S9a**), we utilized TCR transgenic OTII T cells in the IVSA (**Fig. S9b**). OVA peptide was introduced into the culture to mediate MHC II-peptide and OTII TCR interaction so either APCs or MHC II^+^ T_reg_ could stimulate TCR signaling. Overexpression of CIITA in T_reg_ cells led to high levels of MHC II on the cells’ surface (**Fig. S9c-e**) and revealed MHC II^+^ T_reg_ cells were more suppressive than MHC II^low^ T_reg_ cells (**Fig. S9f**).

In the standard IVSA setup, we couldn’t determine whether MHC II-peptide complexes on T_reg_ cells directly interacted with the TCRs of T_eff_ cells (**Fig. 6a**). To overcome this limit, we set up a modified IVSA that used plate coating anti-CD3 and anti-CD28 in place of APCs (**Fig. 6b**). This enabled us to test whether the enhanced suppressive function of MHC II^+^ T_reg_ cells is dependent on antigen-dependent interaction between MHC II^+^ T_reg_ cells and T_eff_ cells by culturing T_reg_ cells and OTII CD4 T_eff_ cells with or without the OVA peptide. The results indicated that CIITA overexpression enhanced T_reg_ function, independent of the presence of the OVA peptide (**Fig. 6c,d**). To further confirm that MHC II is the key functional molecule downstream of CIITA, boosting T_reg_ function, we conducted two loss-of-function assays. In the first, we deleted H2-Ab1, a component of the MHC II, on T_reg_ cells. IVSA revealed that overexpressing CIITA while knocking out MHC II resulted in T_reg_ function returning to normal levels (**Fig. 6e**), indicating that MHC II is indeed the primary functional molecule downstream of CIITA. Moreover, when we administrated an MHC II-blocking antibody in the IVSA, the enhanced T_reg_ function was also reduced to the normal level (**Fig. 6f**). These results suggested that MHC II expression on T_reg_ cells can promote its interaction with TCR of T_eff_ cells in an antigen-independent manner, contributing to increased T_reg_ suppressive function. Given that the TCR is not the only molecule on the cell surface that MHC II can interact with, we hypothesize that MHC II would potentially interact with CD4 and lag3 on T_eff_ cells as well(*49, 50*). Such interaction could strengthen the engagement between T_reg_ and T_eff_ cells, thereby enhancing the suppressive function of T_reg_ cells. The detailed molecular mechanism underlying these interactions is unknown and warrants further investigation.

**Figure 6:**
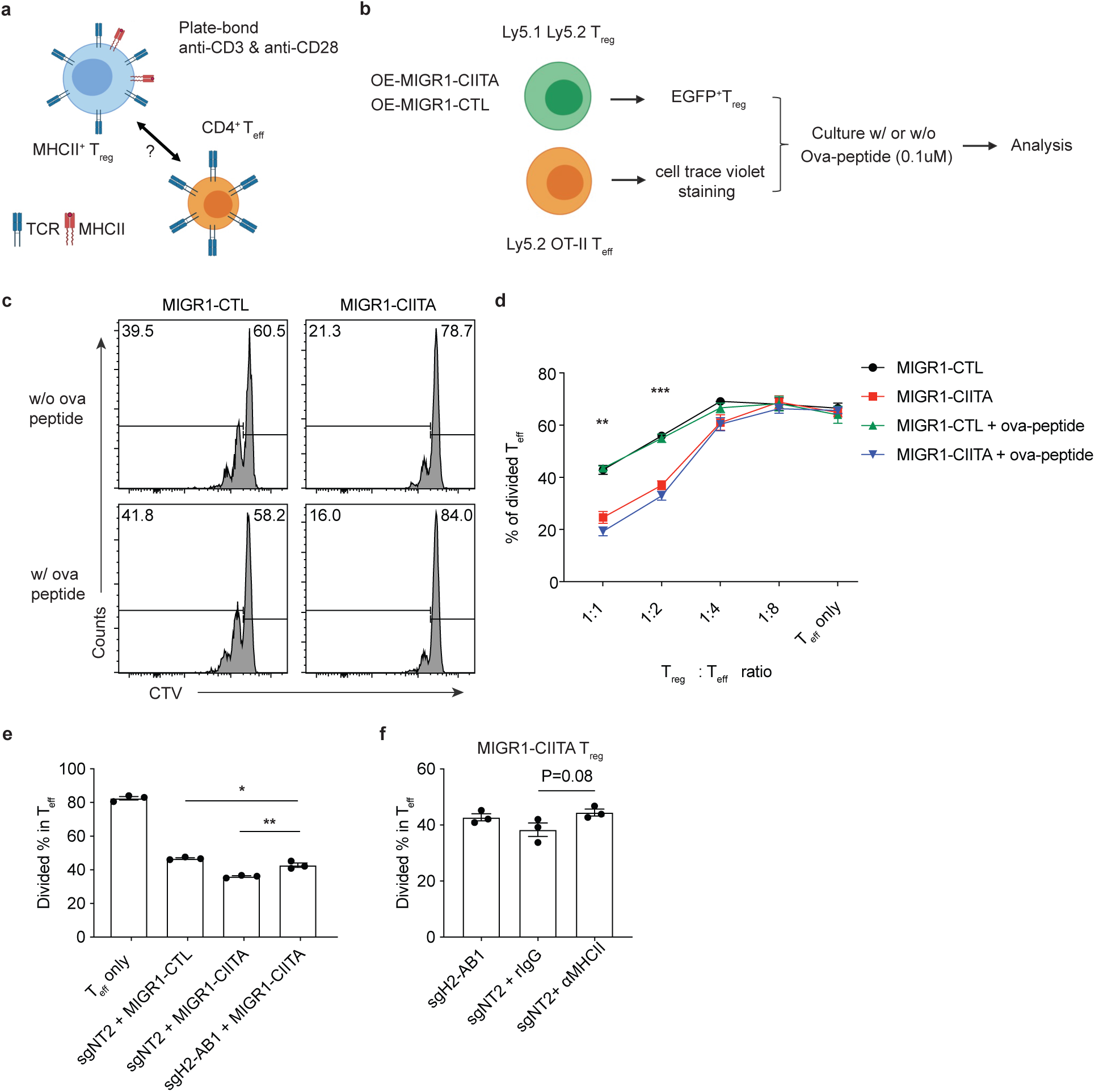
CIITA-MHC II^+^ T_reg_ cells show enhanced suppressive function through two cell type *in vitro* suppression assay. A | Schematic representation of the in vitro suppression assay setup, featuring the interaction between MHC II^+^ T_reg_ cells and CD4^+^ effector T cells (T_eff_). B | Experimental workflow for the T_reg_ cells in vitro suppression assay using the OT-II-Ova peptide system, detailing the pre-coating of a 96-well plate with anti-CD3 and anti-CD28 antibodies prior to T cell culture, and the final concentration of Ova-peptide used. C | Histograms showing T_eff_ cell division within the suppression assay, indicating the functional impact of MHC II+ T_reg_ cells versus control on the proliferative capacity of T_eff_ cells, with and without Ova-peptide. D | Statistical analysis of T_eff_ cell division percentages in the presence of MHC II^+^ T_reg_ cells (MIGR1-CIITA) or control vector (MIGR1-CTL) in the suppression assay. E | Analysis of T_eff_ cell division in assays where T_reg_ cells were engineered with sgNT2 (control) or H2-Ab1 knockout, in the presence of overexpressed MHC II^+^ T_reg_ cells (MIGR1-CIITA), maintaining a T_reg_:T_eff_ ratio of 1:1. F | Division analysis of T_eff_ cells with sgNT2 or H2-Ab1 knockout in MHC II^+^ T_reg_ cells (MIGR1-CIITA) treated with or without an MHC II blocking antibody (αMHC II), in the two cell type in vitro suppression assay system, also with a T_reg_:T_eff_ ratio of 1:1. *Statistical significance is indicated as: *P < 0.05, **P < 0.01, ***P < 0.001. The percentage of divided T_eff_ cells serves as an indicator of T_reg_ suppressive capacity in the assay*.

### MHC II expression on T_reg_ cells strengthens direct interaction between T_eff_ and T_reg_ cells

Recently, the LIPSTIC assay was developed to investigate the direct interactions between cell types via ligand-receptor pairs(*51*). This assay is based on *S. aureus* enzyme Sortase A (SrtA), a transpeptidase that can ligate a substrate peptide LPETG to an N-terminal glycine residue. We utilized this assay to further examine the interactions between MHC II^+^ T_reg_ cells and T_eff_ cells *in vitro*. The ligand-receptor pair of neurexin (NRX-SrtA) and neuroligin (NLG-G5) fused to the SrtA/G5 system were retrovirally expressed on T_reg_ (donor) and T_eff_ (recipient) cells, respectively (**Fig. 7a**). The interaction intensity between T_reg_ and T_eff_ cells was determined by adding biotinylated LPETG peptide substrate and measuring the biotin levels on the T_eff_ “recipient” cell surfaces (**Fig. 7b,c**). The “Donor” T_reg_ cells were also transduced with CIITA to boost MHC II expression. We compared the interaction between donor and recipient cells across two groups: one treated with IgG isotype and the other with anti-MHC II blocking antibody to test the effect of the engagement of MHC II to TCR/CD4/Lag3. Flow cytometry revealed a significantly higher biotin signal on the T_eff_ recipient cell population in the IgG isotype treatment group compared to the anti-MHC II treatment group (**Fig. 7d-h**). This result supported that the expression of MHC II on T_reg_ cells strengthens their interaction with T_eff_ cells, leading to their better immune suppressive function.

**Figure 7:**
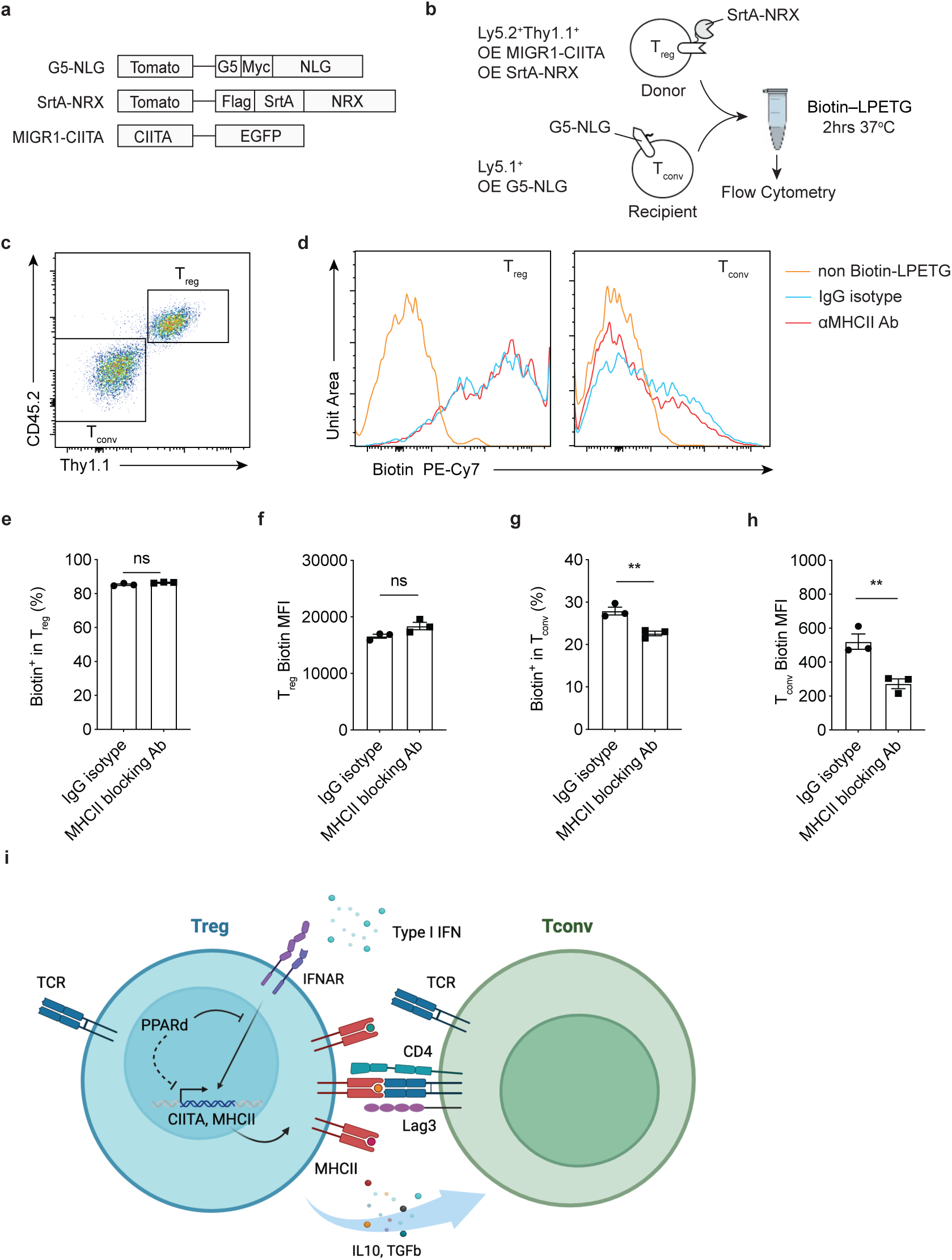
MHC II and TCR/CD4/Lag3 interaction enhance suppressive function of MHC II^+^ T_reg_ cells. A | Diagram illustrating the construction of plasmids used in the LIPSTIC (Labeling of Immune Partnerships by SorTagging Intercellular Contacts) system to investigate intercellular interactions. B | The experimental setup for the LIPSTIC assay to detect physical interactions between T_reg_ and T_eff_. C | Identification of T_reg_ and T_eff_ cell populations using cell surface markers CD45.2 and Thy1.1, respectively, within the LIPSTIC assay. D | Histograms displaying the biotin signal from donor (T_reg_) and recipient (T_eff_) cells post-LIPSTIC assay across different treatment groups. E | Bar graphs quantifying the percentage of biotin-positive T_reg_ cells following the assay. F | Bar graphs presenting the mean fluorescence intensity (MFI) of biotin labeling in T_reg_ cells. G | Bar graphs depicting the percentage of biotin-positive T_eff_ cells after LIPSTIC interaction. H | Bar graphs showing the MFI of biotin labeling in T_eff_ cells, indicating the strength of intercellular interaction. I | Schematic representation of the hypothesized working model based on LIPSTIC assay findings. *Data are presented as mean ± SEM. Statistical significance was evaluated using appropriate statistical tests, with significance indicated as: **P < 0.01, ns denotes not significant*.

## Discussion

Our research has identified the PPARδ/CIITA-MHC II axis as a key regulator of the suppressive function of intra-tumoral T_reg_ cells through MHC II expression. Type 1 IFNs induce the upregulation of CIITA/MHC II in these T_reg_ cells, while PPARδ counteracts this effect by downregulating CIITA/MHC II via suppression of Jak3 expression so that reduces JAK-STAT1 signaling downstream of the IFNα receptor. The presence of MHC II on T_reg_ cells enhances their suppressive function by strengthening the T_reg_- T_eff_ cell interaction, mediated by the engagement of MHC II with TCR/CD4/Lag3 (**Fig. 7i**).

A prior study has reported contradictory results regarding the effects of PPARδ deficiency in T_reg_ cells on tumor growth(*31*). Variations in findings may arise from the different sources of the PPARδ conditional knockout mice used in the two studies (Ppard^tm1Mtz^ vs. Ppard^tm1Rev^), despite targeting the same exon of PPARδ(*43, 52*). Additionally, differences may be introduced by the distinct cancer models employed. Our findings were also substantiated by examining tumor models induced by three different tumor cell lines, including further assessing T_reg_ cell proliferation, apoptosis, and metabolic function. Similar observations on PPARδ’s impact on tumor growth were also made by Dr. Beyaz’s lab in colorectal cancer models. Therefore, PPARδ’s role in curbing intra-tumoral T_reg_ suppression and modulating tumor growth is likely to be relevant to many types of tumors.

It is important to note that human T_reg_ cells include an HLA-DR^+^ population, as MHC II is expressed on activated T cells and serves as an activation marker in human peripheral blood T cells(*53–55*). In contrast to human T cells, MHC II genes in mouse T cells are widely recognized as being silenced(*56*). Our study discovered that the CIITA-MHC II axis was upregulated in PPARδ-deficient mice within the TME. The lower expression level of MHC II in T_reg_ compared to APCs could be attributed to the activation of alternative CIITA promoters(*57*). A compelling question is whether the MHC II^+^ T_reg_ cells we observed are capable of presenting antigens. Studies in humans suggest that activated T cells can express MHC II, enabling antigen presentation, though their effectiveness as antigen-presenting cells is limited by a lack of antigen-capturing ability(*58, 59*). Additionally, a study examining the role of MHC II in human T_reg_ cells demonstrated that the suppressive function of HLA-DR^+^ T_reg_ *in vitro* relies on direct MHC II interactions independent of antigen specificity(*60*). Antibody blocking of MHC II resulted in the loss of T_reg_ suppressive function, which aligns with our findings in mouse MHC II^+^ T_reg_ cells, suggesting a conserved mechanism of MHC II-mediated T_reg_-T_eff_ interaction between mice and humans. Furthermore, structural analyses have shown that specific amino acids in the variable regions CDR1 and CDR2 of TCRs consistently interact with MHC proteins, indicating a basal interaction affinity between TCR and MHC, regardless of the bound antigen(*61*). Together, these findings suggest that T_reg_ cells possess adaptive mechanisms to respond to local environmental cues, potentially through conserved MHC II-mediated interactions that enhance their regulatory function across species.

An increasing number of human studies indicate that DR^+^ T_reg_ cells are correlated with tumor progression and poor prognosis(*62–64*). However, no mechanistic studies have yet elucidated the specific signaling pathways that regulate CIITA-MHC II in human T cells. In our mouse T_reg_ study, we found that PPARδ and type 1 IFN as upstream regulators of the CIITA-MHC II axis. Through RNA-seq and Cut-and-Run analyses, JAK3 emerged as a gene directly regulated by PPARδ, acting as an interception point within the type 1 IFN signaling pathway. JAK3 kinase is critical in cytokine receptor signaling, primarily through its association with the common gamma chain found in cytokine receptors(*65*). Furthermore, it is critical for T cell development, evidenced by the T cell maturation defects in JAK3-deficient mice(*66*). Notably, within the JAK family, JAK1, JAK2, and Tyk2 are expressed at lower levels than JAK3 in tumor-infiltrating T_reg_ cells based on our RNA-seq data, and they do not show differential expression between PPARδ knockout and wild-type cells. We didn’t detect PPARδ binding in their promoter regions. Our finding aligns with prior studies suggesting that, while JAK3 has not been directly observed to phosphorylate STAT1, STAT1 phosphorylation is JAK3-dependent. Disruption of JAK3, whether by knockout or inhibition, leads to decreased STAT1 phosphorylation and subsequent attenuation of type 1 IFN signaling(*67, 68*). In our assays, inhibition of JAK3 by a specific inhibitor reduced the phosphorylation of the STAT1-Y701 site induced by IFN-β. The underlying mechanisms of how JAK3 regulates type 1 IFN signaling warrant further exploration.

Our study identified that PPARδ regulates the CIITA-MHC II axis via JAK3 in T_reg_ cells. It is likely that additional mechanisms may also contribute to this regulation. For instance, PPARδ-mediated modulation of lipid metabolism might influence the epigenetic landscape of the CIITA-MHC II axis in intra-tumoral T_reg_ cells, a possibility that warrants further investigation. Additionally, the limited number of DEGs identified in our RNA-seq analysis and the modest metabolic differences observed between PPARδ WT and KO T_reg_ cells in cellular functional assays may be attributed to compensatory activity by PPARα, a phenomenon previously reported in other cell types(*69*). While our proposed working model highlights type 1 IFN signaling as a key intermediary in the subcutaneous tumor models illustrated, it is important to consider that other signaling pathways may also regulate the CIITA-MHC II axis in different TMEs. These alternative pathways need to be explored in future studies.

In summary, our study uncovers a noncanonical role of PPARδ in restraining the suppressive functions of intra-tumoral T_reg_ cells through the modulation of expression of MHC II. This highlights a critical interplay between intracellular signaling and extracellular environments that convergently shape T_reg_ cell functionality within the TME.

## Methods

### Mice

All mice were maintained in specific pathogen-free facilities at the Salk Institute. Animal experiments were conducted under the regulation of the Institutional Animal Care and Use Committee according to the institutional guidelines. All mice used in the present study are in the C57BL/6 genetic background. C57BL/6 Ly5.1^+^ congenic mice and Rag1^-/-^ mice purchased from the

Jackson Laboratory were used for T_reg_ cell suppression assay and adoptive T cell transfer in B16F10 melanoma models. *Foxp3*^YFP-Cre^*PPARδ^fl/fl^* mice were generated by crossing *Foxp3*^YFP-cre^ mice26 with *PPARδ*^flox^ mice (Jackson laboratory Strain #: 005897). C57BL/6 Rosa-Cas9/Foxp3Thy1.1 mice were generated by crossing Rosa26-LSL-Cas9 mice (The Jackson Laboratory #024857) with *Foxp3*^Thy1.1^ reporter mice. *Foxp3*^Thy1.1^ reporter mice were used to isolate T_reg_ cells for over-expression CIITA in *in vitro* suppression assay and *Foxp3*^Thy1.1^ reporter mice or Rosa-Cas9/ *Foxp3*^Thy1.1^ were used to isolate T_reg_ cells for adoptive transfer assay to validate the function of PPARδ and CIITA upstream of MHC II. Thymus T cell differentiation analysis was checked when mice were around 6 weeks old. All other experiments were initiated in the 8- to 10-week-old male or female mice, unless otherwise specified. All mice used in experiments were socially housed under a 12 h light: dark cycle, with an ambient temperature of 20–26 °C and humidity of 30–70%.

### In vitro culture of T_reg_ cells

IL2 expanded T_reg_ cells (ref of IL2 expansion) were isolated from the spleen and peripheral lymph nodes of Foxp3^Thy1.1^ reporter mice or Rosa-Cas9 *Foxp3*^Thy1.1^ mice by anti-PE magnetic beads (Miltenyi, catalog no. 130-048-801) for Cut & Run and adoptive transfer experiment.

### In vitro suppression assay

T_reg_ cells were transduced by retrovirus expressing sgRNA targeting gene of interest or retrovirus overexpressing CIITA gene. T_reg_ cells were cultured in RPMI complete media supplemented with IL-2 (500 U/ml). Four days after transduction, transduced cells were sorted and mixed with FACS-sorted CD45.1^+^ naive CD4 T cells (CD4^+^ CD25^−^ CD44^lo^ CD62L^hi^) labeled with CellTrace Violet (Thermo Fisher Scientific #C34571) in different ratios in the presence of irradiated T cell depleted spleen cells as antigen-presenting cells (APC). Three days later, T_reg_ suppression function was measured by the percentage of non-dividing cells within the CD45.1^+^ T_eff_ cell population. For two cell-type IVSA experiments, plate-bound anti-CD3 and anti-CD28 antibodies were used to replace APCs. For specific antigen-mediated cell-cell interaction assay, T_eff_ cells were derived from OTII mice, and T_reg_ cells were derived from Thy1.1 reporter mice or Cas9-Thy1.1 reporter mice. T_reg_ suppression readout was measured after three days of co-culture.

### Retroviral production and T cell transduction

HEK293T cells were seeded in 6-wells plate at 0.5 million cells per 2mL DMEM media supplemented by 10% FBS, 1% Pen/Strep, 1 × GlutaMax, 1 × Sodium Pyruvate, 1 × HEPES, and 55 mM beta-mercaptoethanol. One day later, cells from each well were transfected with 1.2 μg of targeting vector pSIRG-NGFR(*70*) or pMIGR1 (for overexpress CIITA) and 0.8 μg of packaging vector pCL-Eco (Addgene, #12371) by using Lipofectamine 3000 (Thermo Fisher, #L3000008) according to manufactured protocol. Cell culture media was replaced by 2mL fresh DMEM complete media at 24 hours and 48 hours after transfection. The retroviral supernatant was collected at 48 and 72 hours post-transfection for T cell infection. For experiments with CRISPR sgRNA targeting, Cas9^+^ T_reg_ cells were first seeded in 24-well plates coated with anti-CD3 and anti-CD28 antibodies. At 24 hours post-activation, T_reg_ media from each well was replaced by retroviral supernatant, supplemented with 4 μg/mL Polybrene (Millipore # TR-1003-G), and spun in a benchtop centrifuge at 1,258 x g for 90 minutes at 32°C. After centrifugation, T_reg_ media was replaced with fresh media supplemented with human IL-2 and cultured for another three days. Transduced cells were analyzed for Foxp3 and cytokine expression in eBioscience Fix/Perm buffer (eBioscience #00-5523-00) using flow cytometry. Transduced NGFR^+^ cells were FACS-sorted for subsequent *in vitro* adoptive transfer assay and Cut and Run experiments.

### Cut-and-Run

We adopt the same procedure of Cut & Run for T_reg_ cells(*71*), which is modified from the original Cut & Run protocol(*72*).

### RNA isolation, RNA-seq, and RT–qPCR

RNA was isolated using TRIzol RNA isolation reagent (Invitrogen). RNA concentration and integrity were determined by Bioanalyzer using RNA 6000 Pico Kit (Agilent). RNA-seq libraries were prepared using Illumina TruSeq Stranded mRNA kit (Illumina) following the manufacturer’s instructions. Complementary DNA was synthesized using SuperScript IV First-Strand Synthesis System (Thermo Fisher Scientific, catalog no. 18091050). RT–qPCR was performed using Power SYBR Green Master Mix (Thermo Fisher Scientific, catalog no. 4309155) on a ViiA 7 Real-Time PCR System. The relative quantification value was calculated as 2^−ΔCt^ relative to internal control (*Hprt*). Details of primer sequences are listed in the Supplementary Table 1.

### Adoptive T cells transfer

T_reg_ cells were purified from the spleens and lymph nodes of IL2-expanded mice, and transduced by retrovirus expressing sgRNA targeting gene of interest, and cultured in RPMI complete media and IL-2 (500 U/ml). Four days after transduction, the NGFR^+^ transduced T_reg_ cells were FACS sorted before being transferred into recipient mice. Alternatively, T_reg_ cells were electroporated by CRISPR-sgRNA RNP. T_reg_ cells were co-transferred into *Rag1*^-/-^ recipient mice with T_eff_ cells (purified by anti-PE magnetic beads system and followed by CD3 T cell isolation, Biolegend # 480024)

### Tumor models

*Foxp3*^YFP-Cre^, *PPARδ*^fl/fl^*Foxp3*^YFP-Cre^ mice were injected with B16.F10 melanoma (2.5× 10^5^ cells intradermally), MC38 colon carcinoma (5 × 10^5^ cells subcutaneously), EL4 thymoma (5 × 10^5^ cells intradermally). Mice were randomized co-housing before tumor implantation. Tumors were measured regularly with digital calipers and tumor volumes were calculated; this was done blindly. Tumors and spleens were collected for analysis. TILs were prepared using a 47% Percoll gradient followed by mechanical disruption and collagenase (TL collagenase, Roche #05401020001), DNase I (Roche #4716728001) digestion, and passed through 100 μm cell strainer to collect single cell suspension. Isolated cells were stimulated with PMA/Ionomycin and Golgi plug for 5 hours, and then were subjected to Foxp3 and cytokines staining with eBioscience Fix/Perm buffer (eBioscience #00-5523-00). For T cell adoptive transfer tumor models, B16.F10 tumor cells were implanted into *Rag1*^-/-^ recipient mice three days post T cell transfer.

### RNP electroporation

Fresh isolated T_reg_ cells and T_eff_ cells were subjected to CRISPR/Cas9 knockout by Lonza 4D-NulceofecorTM system and P3 primary cell 4D Nucleofector electroporation kit (Lonza, Cat# V4XP-3032 for electroporation wells) according to the manufacture protocols. 40 pmol Recombinant Cas9 protein (Integrated DNA Technologies (IDT), Cat#1081059) and 150 pmol 20bp sgRNAs (Synthego, CRISPR-evolution sgRNA EZ Kit). Electroporated T cells were recovered for 20 minutes before *in vivo* adoptive transfer.

## Acknowledgements

We extend our heartfelt gratitude to all members of the Zheng lab for their invaluable assistance and insightful suggestions throughout this work. We also thank the Salk Razavi Newman Integrative Genomics and Bioinformatics Core for their expert support with sequencing data analysis. We would like to thank Matthew Maxwell, Thomas Mann, Alexandra G. Moyzis, and Kay Chun at the Salk NOMIS Center for their suggestions and assistance. Q.Y. was supported by a NOMIS Fellowship. J.Y. was supported by the National Institutes of Health (NCI P30-CA014195, NIA P01-AG073084, NIA-NMG RF1-AG064049, NIA P30-AG068635). Y.Z. was supported by the NOMIS Foundation, the Sol Goldman Trust, and the National Institutes of Health (R01-AI107027, R01-AI1511123, R21-AI178938, S10-OD023689, and S10-OD034268). This study was also supported by National Cancer Institute funded Salk Institute Cancer Center Core Facilities (P30-CA014195).

## Supplementary Figure legends

**Supplementary Figure 1:**
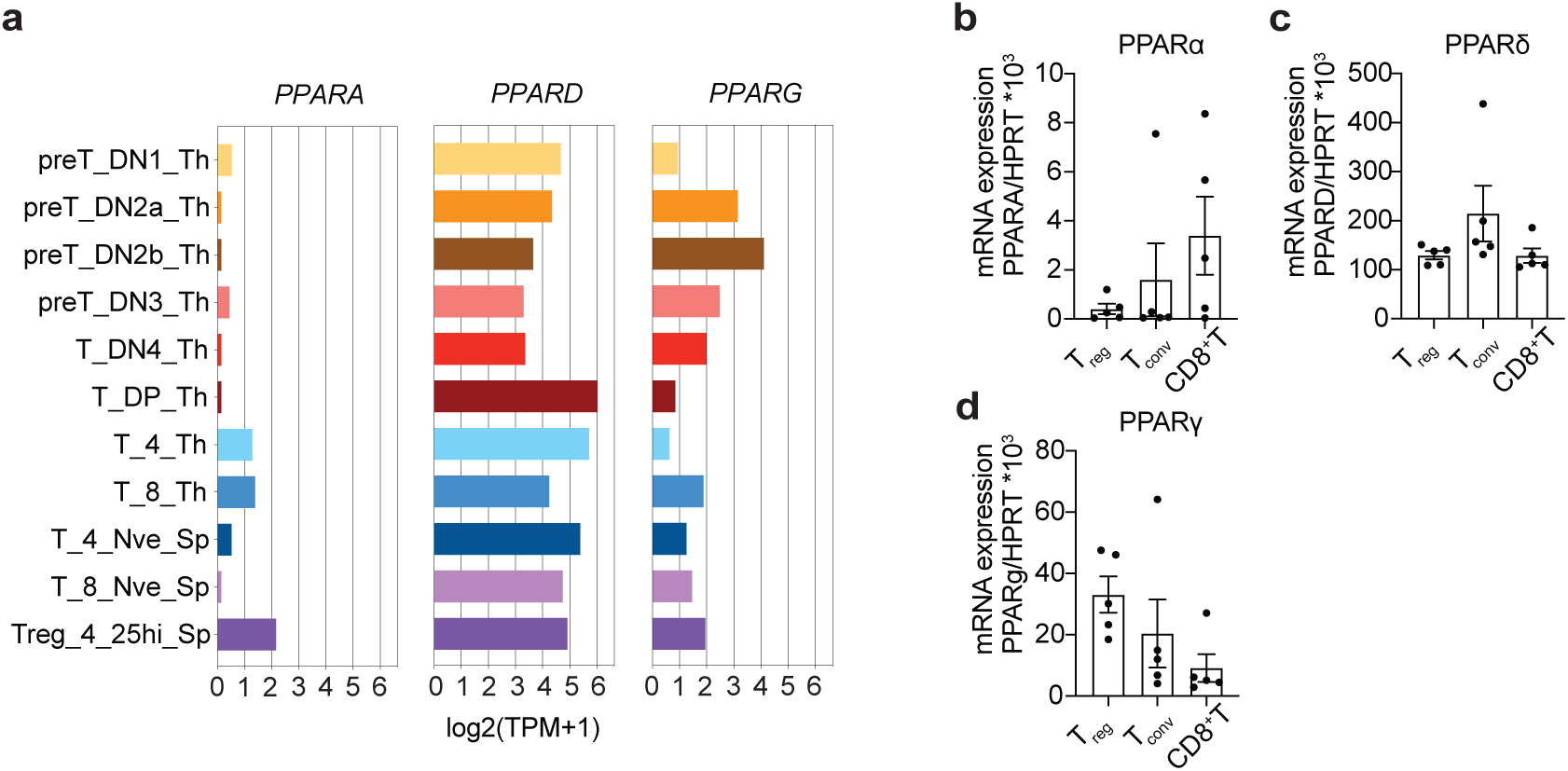
Differential Expression of PPAR Isoforms in T Cell Subsets. A | Expression levels of PPAR family genes (PPARα, PPARδ, and PPARγ) across various stages of T cell differentiation, utilizing the ULI-RNAseq database from the ImmGen project. B-D | Quantitative RT-PCR analysis of PPAR isoform gene expression in sorted, purified naive T_reg_, conventional T cells (T_conv_), and CD8^+^ T cells from C57BL/6 wild-type mice, aged 8-12 weeks (n=5). Relative expression levels are shown for each PPAR isoform within the different T cell populations. *Data are represented as mean values ± SEM*.

**Supplementary Figure 2:**
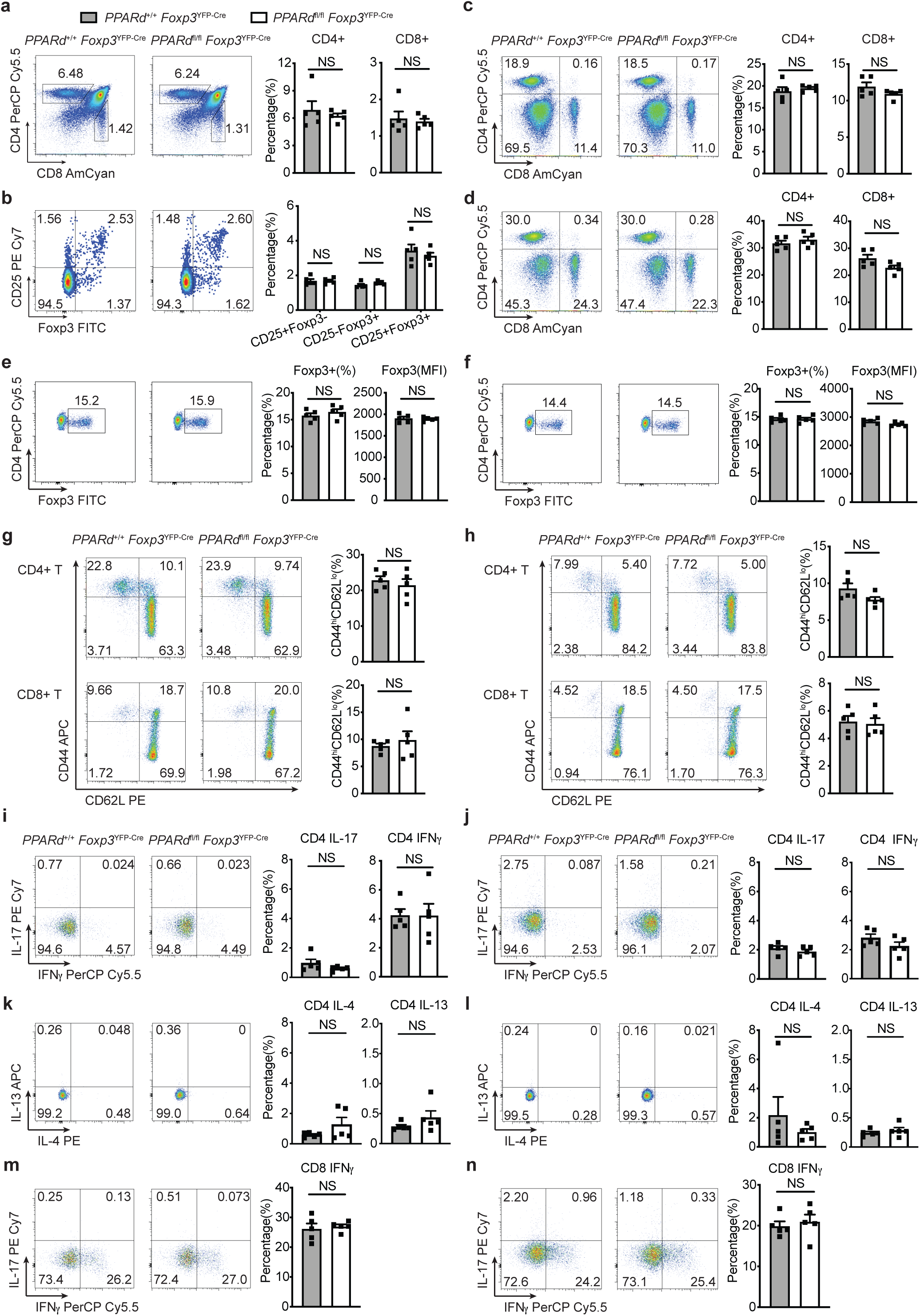
Normal T_reg_ development and function in PPARδ cKO mice under steady-state conditions. A | Flow cytometric analysis comparing the percentage of CD4^+^ and CD8^+^ T cells in the thymus of wild-type (WT) and PPARδ conditional knockout (cKO) mice. B | Quantification of CD25^+^Foxp3^−^, CD25^−^Foxp3^+^, and CD25^+^Foxp3^+^ T_reg_ progenitor and mature T_reg_ cells in the thymus of WT and PPARδ cKO mice. C, D | Flow cytometric analysis of CD4^+^ and CD8^+^ T cell populations in the spleen (C) and peripheral lymph nodes (D) of WT and PPARδ cKO mice. E, F | Percentage and mean fluorescence intensity (MFI) of Foxp3^+^ T_reg_ cells in the spleen (E) and peripheral lymph nodes (F). G, H | Analysis of activated/memory CD4^+^ and CD8^+^ T cells, characterized as CD44^high^CD62L^low^, in the spleen (G) and peripheral lymph nodes (H). I, J | Proportion of IFNγ or IL-17 producing CD4^+^ T cells in the spleen (I) and peripheral lymph nodes (J). K, L | Frequency of IL-4 or IL-13 producing CD4^+^ T cells in the spleen (K) and peripheral lymph nodes (L). M, N | Quantification of IFNγ producing CD8^+^ T cells in the spleen (M) and peripheral lymph nodes (N). *Statistical significance was assessed using a two-tailed unpaired Student’s t-test, with no significant difference (NS) observed in the measured parameters between the WT and PPARδ cKO groups. Data are represented as mean ± SEM*.

**Supplementary Figure 3:**
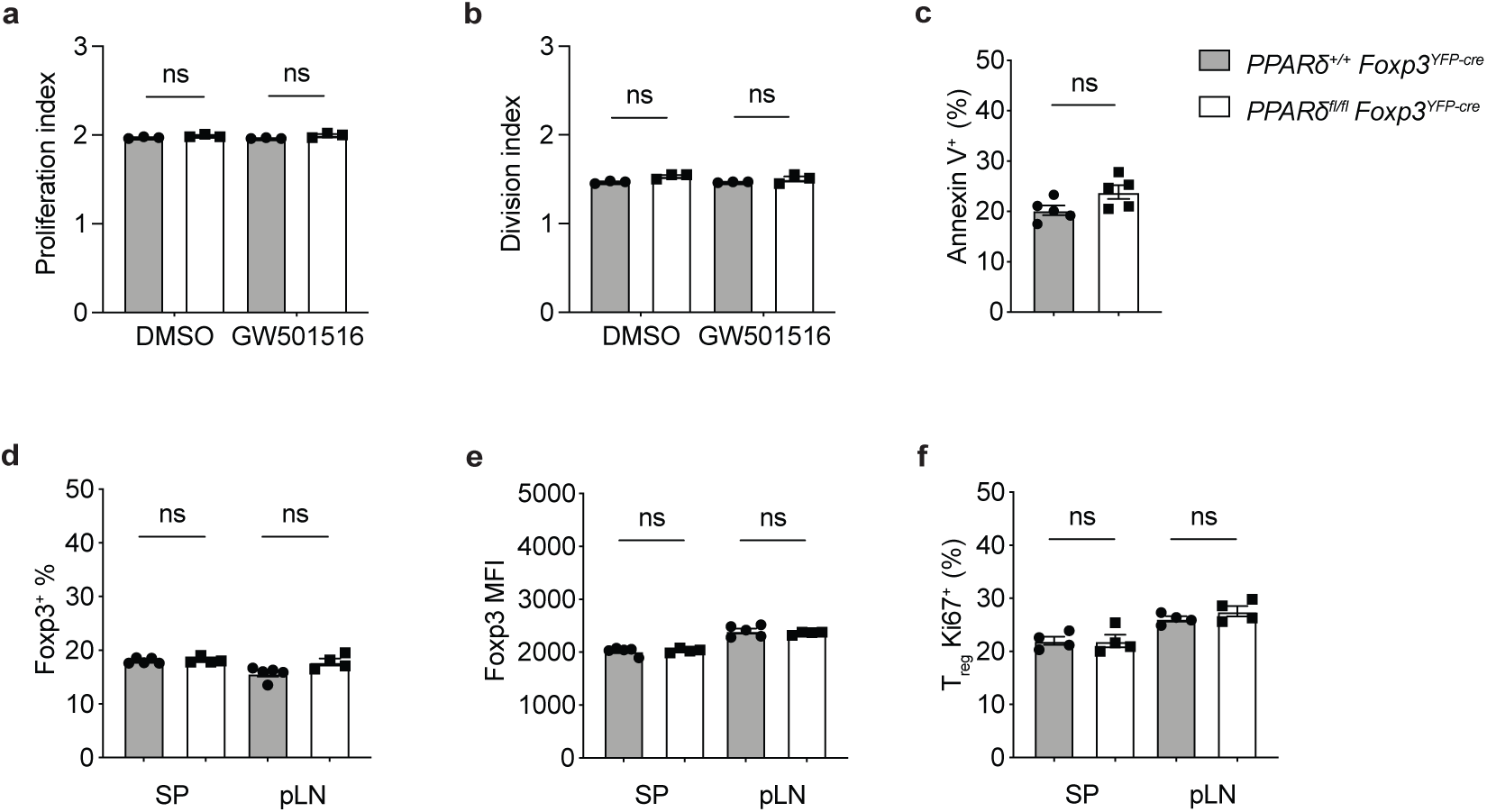
Proliferation, apoptosis, and homeostasis of Foxp3^cre^PPARδ^fl/fl^ T_reg_ cells. A | The proliferation index of nature T_reg_ cells (nT_reg_) from PPARδ^+/+^ and PPARδ^fl/fl^ mice, determined by Cell Trace Violet dilution on day 3 post-staining, with data analyzed by FlowJo software. B | The division index for the same nT_reg_ populations as in (A), calculated to assess cell divisions over the same period. C | Analysis of apoptosis levels in T_reg_ cells, assessed by annexin V staining of overnight-cultured splenocytes from PPARδ^+/+^ and PPARδ^fl/fl^ mice. D | Percentage of Foxp3^+^ T_reg_ cells in the spleen (SP) and peripheral lymph nodes (pLN) of PPARδ^+/+^ and PPARδ^fl/fl^ mice. E | Mean fluorescence intensity (MFI) of Foxp3 expression in T_reg_ cells from the spleen and peripheral lymph nodes. F | In vivo proliferation of T_reg_ cells evaluated by Ki67 staining in the spleen and peripheral lymph nodes. (n=4) *Each analysis used biological triplicates or a sample size of n=5 mice. No significant differences (ns) were observed between the groups*.

**Supplementary Figure 4:**
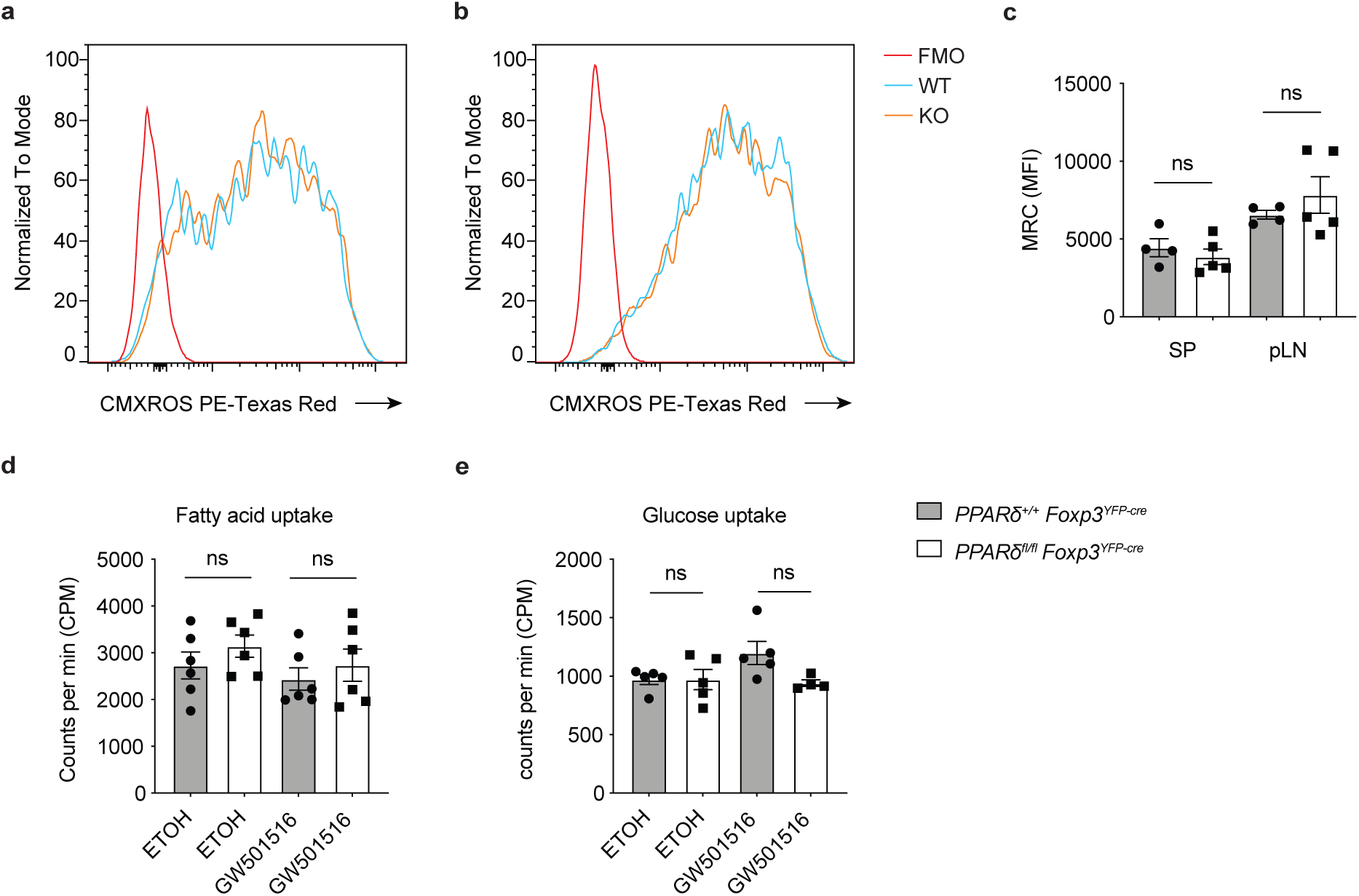
Mitochondrial function and nutrient uptake in Foxp3^cre^PPARδ^fl/fl^ T_reg_ cells. A | Histograms depicting the mitochondrial membrane potential in T regulatory cells (T_reg_) from the spleen and peripheral lymph nodes (pLN), assessed using CMXRos staining. Fluorescence intensities represent mitochondrial potential relative to fluorescence minus one (FMO) controls, and comparisons are made between wild-type (WT) and PPARδ knockout (KO) T_reg_ cells. B | The mean fluorescence intensity (MFI) of mitotracker red CMXRos in T_reg_ cells from the spleen and pLN, comparing WT and KO cells to assess mitochondrial activity. C | Quantitative representation of fatty acid uptake in T_reg_ cells, indicated by counts per minute (CPM), comparing cells treated with ethanol (EtOH) as control and those treated with the PPARδ agonist, GW501516. D | Glucose uptake assay results, also shown as CPM, in T_reg_ cells treated with EtOH or GW501516, across WT and KO groups to examine metabolic function. Data are expressed as mean ± SEM. Statistical significance was assessed based on data distribution and variance characteristics, with ‘ns’ indicating not significant (p > 0.05).

**Supplementary Figure 5:**
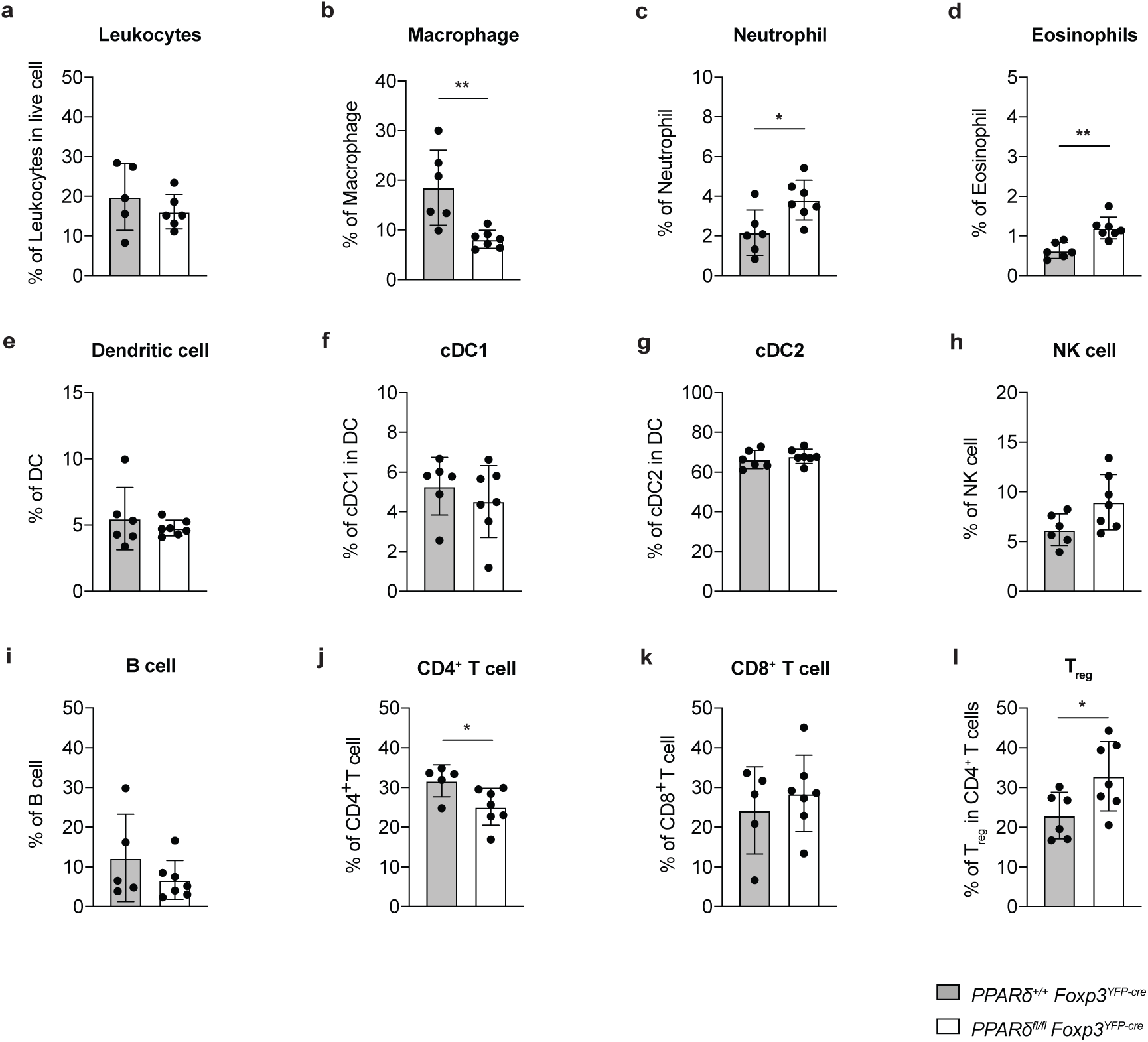
Immune cell profiling in B16F10 melanoma of PPARδ Knockout and Wild-Type Mice. This figure illustrates the immune cell distribution within the tumor microenvironment (TME) of B16F10 melanoma-bearing mice, with a comparison between mice harboring a PPARδ knockout in Foxp3-expressing cells (*PPARδ^fl/fl^Foxp3^YFP-cre^*) and wild-type (*PPARδ^+/+^Foxp3^YFP-cre^*) mice. Immune cells were characterized and quantified as follows: A | Leukocytes identified as CD45.2^+^ cells. B | Macrophages characterized by F4/80+CD11b^+^ markers. C | Neutrophils represented by CD11b^+^Ly-6G^+^. D | Eosinophils designated as CD11b^+^siglec-F^+^. E | Dendritic cells classified by CD11c^+^MHC II^+^ expression. F | cDC1 subset within dendritic cells. G | cDC2 subset within dendritic cells. H | NK cells classified by NK1.1^+^. I | B cells identified as CD19^+^. J | CD4^+^ T cells recognized by TCRb^+^CD4^+^ markers. K | CD8^+^ T cells marked by TCRb^+^CD8^+^. L | T regulatory cells (T_reg_) are defined as TCRb^+^CD4^+^Foxp3^+^ within Ghost-dye^-^CD4^+^ T cells. *Data are presented as mean ± SEM. The percentage of each immune cell type is reported relative to the total immune cell population. Statistical significance was evaluated using a two-tailed unpaired t-test, with significance denoted by *P < 0.05 and **P < 0.01*.

**Supplementary Figure 6:**
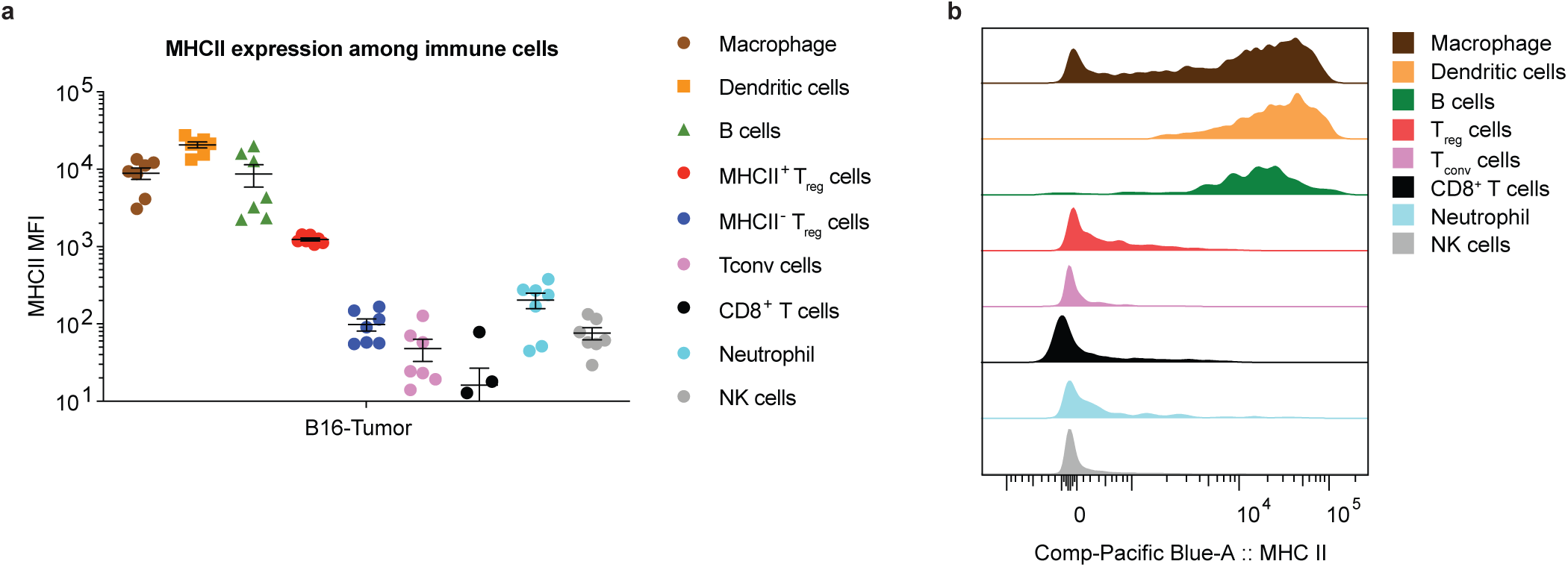
MHC II Expression Across Immune Cell Populations in B16F10 Tumor-Bearing Mice with PPARδ Deficiency. A | Mean fluorescence intensity (MFI) of MHC II expression across various immune cell types within the tumor microenvironment of B16F10 melanoma-bearing mice. This panel compares MHC II levels in cells from PPARδ^+/+^Foxp3^YFP-cre^ (wild-type) and PPARδ^fl/fl^Foxp3^YFP-cre^ (PPARδ-deficient) mice. B | Histogram illustration of MHC II expression presented as MFI for macrophages, dendritic cells, B cells, T regulatory (T_reg_) cells, conventional T cells (T_conv_), CD8^+^ T cells, neutrophils, and natural killer (NK) cells. *Immune cells were gated based on their specific surface markers and analyzed for MHC II expression using flow cytometry. MFI data are presented on a logarithmic scale to allow for comparison across different cell types*.

**Supplementary Figure 7:**
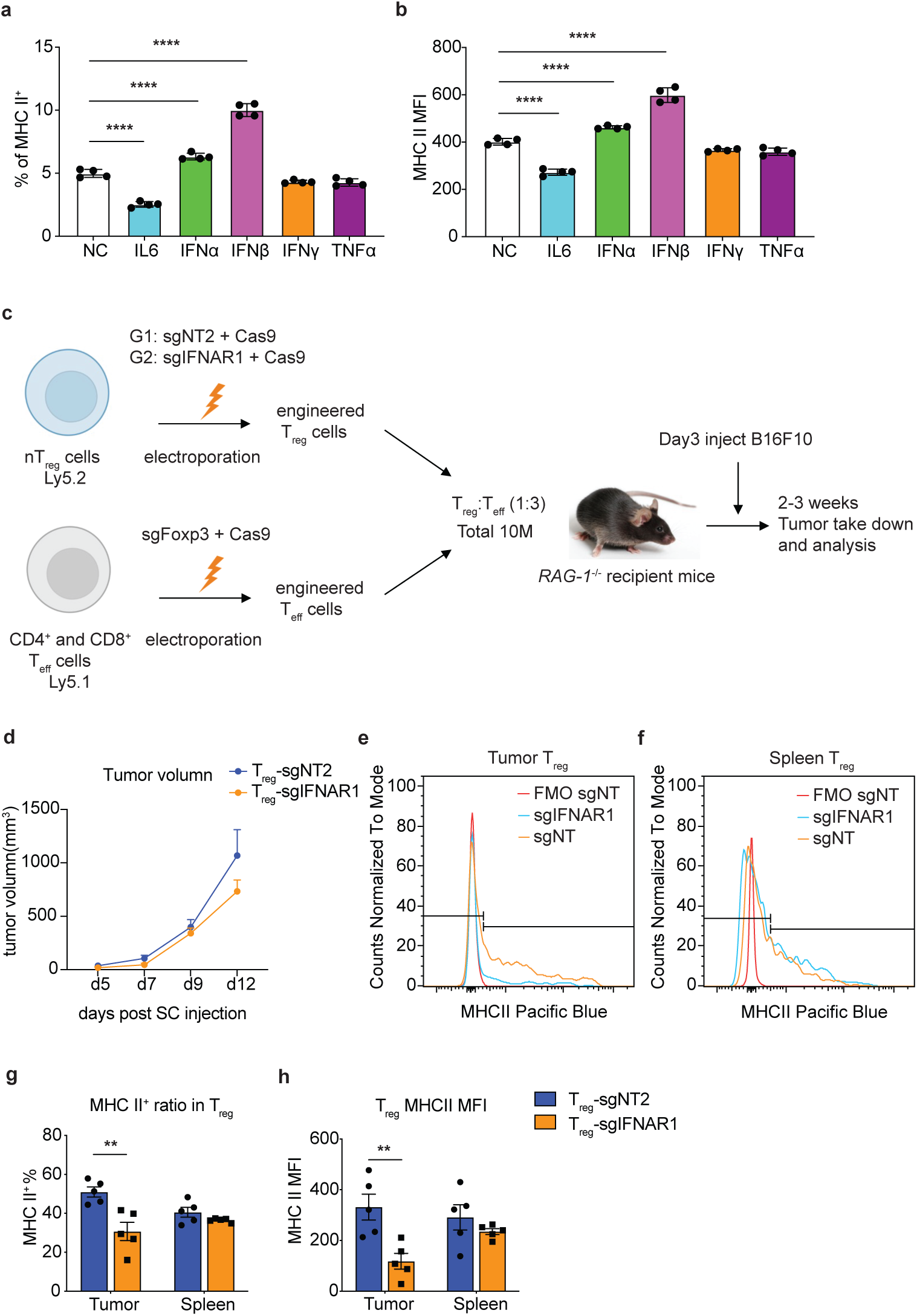
Influence of Cytokines on MHC II Expression in T_reg_ cells and Functional Analysis of IFNAR1-Deficient T_reg_ cells in Tumor Context. A | In vitro analysis of MHC II expression induction in T_reg_ cells after treatment with a panel of cytokines, including Negative control (NC), mIL-6, mIFN-α, mIFN-β, mIFN-γ, and hTNFα. B | Mean fluorescence intensity (MFI) of MHC II on T_reg_ cells following cytokine treatments as compared to the negative control). C | Schematic representation of the adoptive transfer model used to investigate the function of T_reg_ cells deficient in IFNAR1 and wild-type (WT) T_reg_ cells in a B16F10 melanoma tumor model. D | Tumor volume measurements over time following subcutaneous injection of B16F10 cells in RAG-1-/- recipient mice that were adoptively transferred with engineered T_reg_ cells, either with sgNT2 (non-targeting control) or sgIFNAR1. E | Histograms showing the expression of MHC II on T_reg_ cells isolated from tumor tissue, comparing T_reg_ cells with sgNT2 and sgIFNAR1. F | Histograms of MHC II expression on T_reg_ cells from the spleen, contrasting sgNT2 and sgIFNAR1 conditions. G | Bar graphs quantifying the percentage of MHC II^+^ T_reg_ cells in both the tumor and spleen across the sgNT2 and sgIFNAR1 groups. H | Bar graphs presenting the MFI of MHC II on T_reg_ cells, comparing tumor-infiltrating and splenic T_reg_ cells following the adoptive transfer of T_reg_ cells with sgNT2 and sgIFNAR1. *Statistical significance was determined using two-tailed P values with unpaired t-tests, with **P < 0.01*.

**Supplementary Figure 8:**
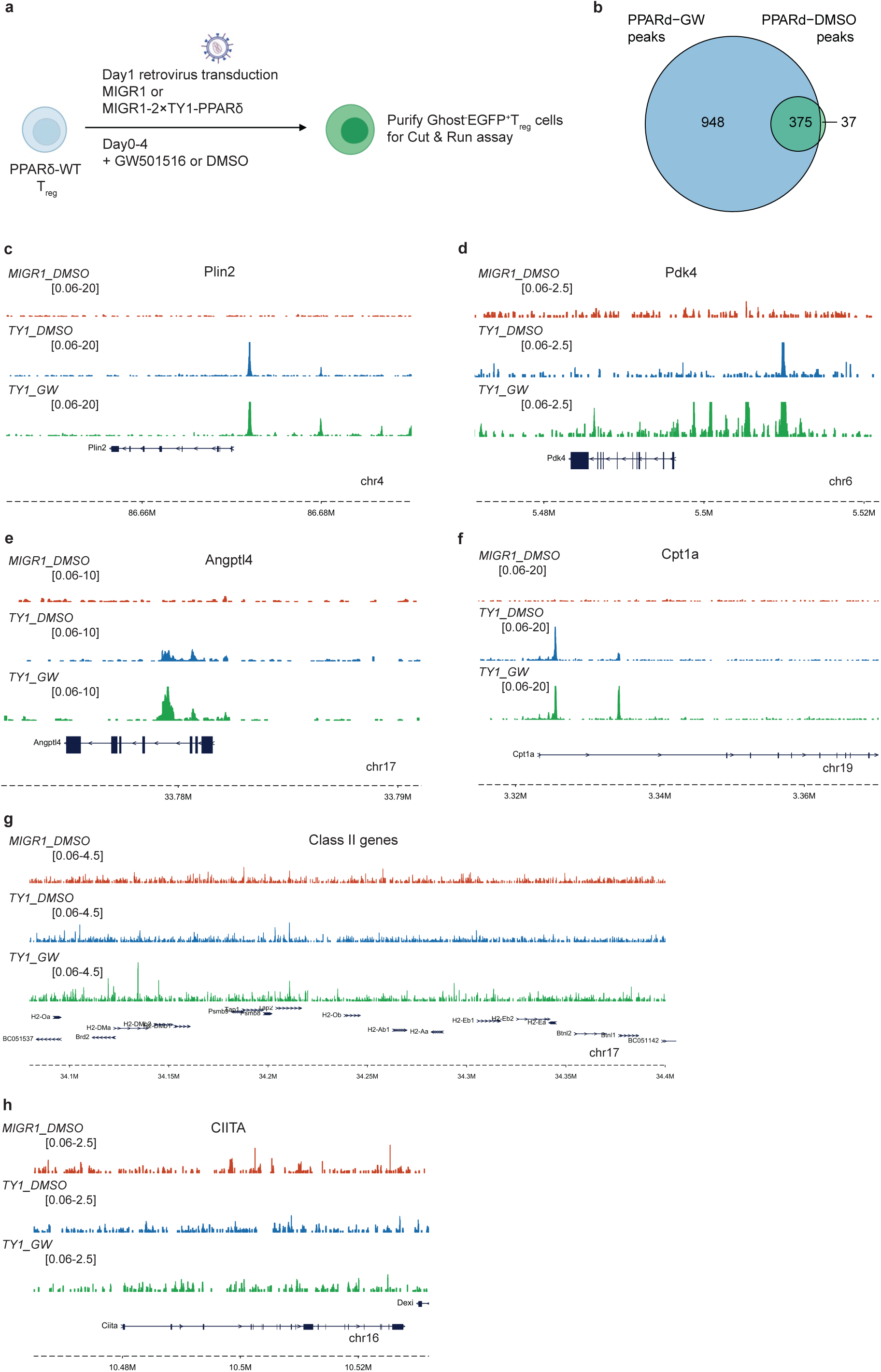
Cut & Run Analysis of PPARδ and Foxp3 Binding Sites in Cultured T_reg_ cells. A-C | A | Experimental setup for retroviral transduction of T_reg_ cells with MIGR1 vector expressing TY1-PPARδ. T_reg_ cells were treated with the PPARδ agonist GW501516 or DMSO. B | Venn diagrams illustrating the overlap of binding peaks of overexpressing MIGR1-TY1-PPARδ T_reg_ cells between PPARδ agonist (GW) treatment and vehicle (DMSO) treatment. C-H | Genomic tracks showcasing peaks at different gene loci: C | Plin2 locus showing binding peaks with different treatments. D | Pdk4 locus with displayed peaks. E | Angptl4 locus peaks in the context of different conditions. F | Cpt1a locus and its binding patterns under various treatments. G | Representation of class II gene loci without any significant PPARδ binding. H | CIITA locus indicating the absence of PPARδ binding peaks across all conditions.

**Supplementary Figure S9:**
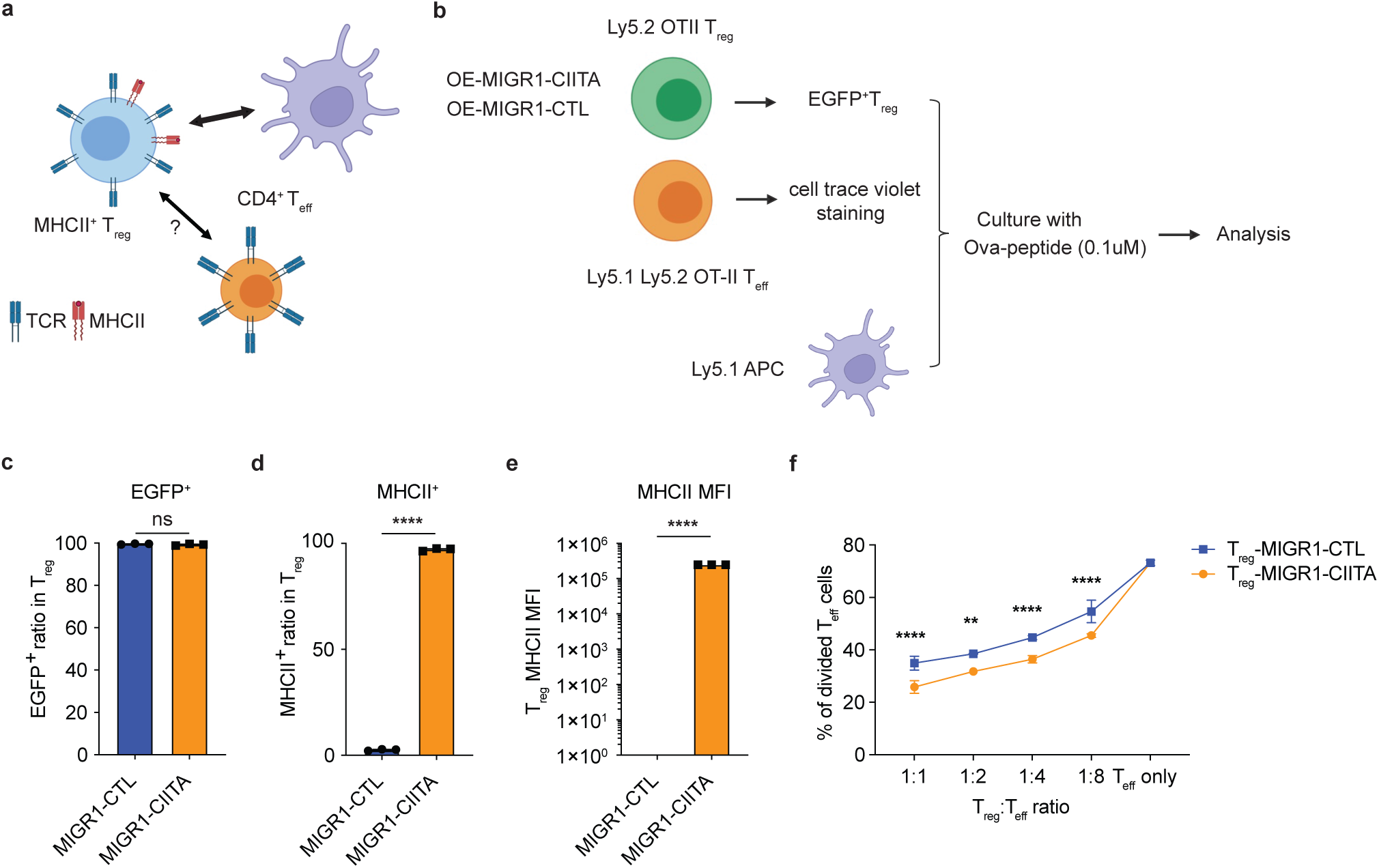
Functional Assessment of CIITA-Overexpressing MHC II^+^ T_reg_ cells Using an In Vitro Suppression Assay. A | Schematic representation of the in vitro T_reg_ suppression assay designed to evaluate the suppressive capability of T_reg_ cells. B | Detailed experimental setup of the suppression assay using the OT-II Ova peptide system, including Tregs overexpressing MIGR1-CIITA or control vector (MIGR1-CTL), T effector cells (T_eff_) and Antigen-presenting cells (APC), cultured with a final concentration of 0.1μM Ova-peptide. C | Proportion of EGFP^+^ T_reg_ cells in the culture, indicating transduction efficiency. D | Percentage of MHC II^+^ cells within the EGFP^+^ T_reg_ population, assessing the upregulation of MHC II due to CIITA overexpression. E | Mean fluorescence intensity (MFI) of MHC II expression on T_reg_ cells, comparing the impact of MIGR1-CIITA to the control. F | Quantification of T_eff_ cell division within the suppression assay, measured at different T_reg_:T_eff_ ratios, illustrating the enhanced suppressive function of CIITA-overexpressing T_reg_ cells. *Statistical significance was determined using a two-tailed unpaired t-test with indicated p-values (**P < 0.01; ****P < 0.0001). The data, comprising three biological replicates per group, are presented as mean values ± SD*.

## Reference

1. A. Tanaka, S. Sakaguchi, Regulatory T cells in cancer immunotherapy. Cell Res 27, 109–118 (2017).

2. G. J. Bates et al., Quantification of regulatory T cells enables the identification of high-risk breast cancer patients and those at risk of late relapse. J Clin Oncol 24, 5373–5380 (2006).

3. T. Sasada, M. Kimura, Y. Yoshida, M. Kanai, A. Takabayashi, CD4+CD25+ regulatory T cells in patients with gastrointestinal malignancies: possible involvement of regulatory T cells in disease progression. Cancer 98, 1089–1099 (2003).

4. T. J. Curiel et al., Specific recruitment of regulatory T cells in ovarian carcinoma fosters immune privilege and predicts reduced survival. Nat Med 10, 942–949 (2004).

5. E. Sato et al., Intraepithelial CD8+ tumor-infiltrating lymphocytes and a high CD8+/regulatory T cell ratio are associated with favorable prognosis in ovarian cancer. Proc Natl Acad Sci U S A 102, 18538–18543 (2005).

6. M. W. Teng et al., Multiple antitumor mechanisms downstream of prophylactic regulatory T-cell depletion. Cancer Res 70, 2665–2674 (2010).

7. E. Pastille et al., Transient ablation of regulatory T cells improves antitumor immunity in colitis-associated colon cancer. Cancer Res 74, 4258–4269 (2014).

8. G. Q. Phan et al., Cancer regression and autoimmunity induced by cytotoxic T lymphocyte-associated antigen 4 blockade in patients with metastatic melanoma. Proc Natl Acad Sci U S A 100, 8372–8377 (2003).

9. S. Sakaguchi, T. Yamaguchi, T. Nomura, M. Ono, Regulatory T cells and immune tolerance. Cell 133, 775–787 (2008).

10. N. S. Joshi et al., Regulatory T Cells in Tumor-Associated Tertiary Lymphoid Structures Suppress Anti-tumor T Cell Responses. Immunity 43, 579–590 (2015).

11. C. Liu, C. J. Workman, D. A. Vignali, Targeting regulatory T cells in tumors. FEBS J 283, 2731–2748 (2016).

12. P. Attia et al., Autoimmunity correlates with tumor regression in patients with metastatic melanoma treated with anti-cytotoxic T-lymphocyte antigen-4. J Clin Oncol 23, 6043–6053 (2005).

13. O. Annacker, R. Pimenta-Araujo, O. Burlen-Defranoux, A. Bandeira, On the ontogeny and physiology of regulatory T cells. Immunol Rev 182, 5–17 (2001).

14. Y. Belkaid, C. A. Piccirillo, S. Mendez, E. M. Shevach, D. L. Sacks, CD4+CD25+ regulatory T cells control Leishmania major persistence and immunity. Nature 420, 502–507 (2002).

15. M. J. McGeachy, L. A. Stephens, S. M. Anderton, Natural recovery and protection from autoimmune encephalomyelitis: contribution of CD4+CD25+ regulatory cells within the central nervous system. J Immunol 175, 3025–3032 (2005).

16. T. Takahashi et al., Immunologic self-tolerance maintained by CD25+CD4+ naturally anergic and suppressive T cells: induction of autoimmune disease by breaking their anergic/suppressive state. Int Immunol 10, 1969–1980 (1998).

17. M. Tekguc, J. B. Wing, M. Osaki, J. Long, S. Sakaguchi, Treg-expressed CTLA-4 depletes CD80/CD86 by trogocytosis, releasing free PD-L1 on antigen-presenting cells. Proc Natl Acad Sci U S A 118, (2021).

18. X. Cao et al., Granzyme B and perforin are important for regulatory T cell-mediated suppression of tumor clearance. Immunity 27, 635–646 (2007).

19. D. C. Gondek, L. F. Lu, S. A. Quezada, S. Sakaguchi, R. J. Noelle, Cutting edge: contact-mediated suppression by CD4+CD25+ regulatory cells involves a granzyme B-dependent, perforin-independent mechanism. J Immunol 174, 1783–1786 (2005).

20. T. Bopp et al., Cyclic adenosine monophosphate is a key component of regulatory T cell-mediated suppression. J Exp Med 204, 1303–1310 (2007).

21. S. Deaglio et al., Adenosine generation catalyzed by CD39 and CD73 expressed on regulatory T cells mediates immune suppression. J Exp Med 204, 1257–1265 (2007).

22. S. Paust, L. Lu, N. McCarty, H. Cantor, Engagement of B7 on effector T cells by regulatory T cells prevents autoimmune disease. Proc Natl Acad Sci U S A 101, 10398–10403 (2004).

23. S. Sakaguchi, K. Wing, Y. Onishi, P. Prieto-Martin, T. Yamaguchi, Regulatory T cells: how do they suppress immune responses? Int Immunol 21, 1105–1111 (2009).

24. C. Raffin, L. T. Vo, J. A. Bluestone, Treg cell-based therapies: challenges and perspectives. Nat Rev Immunol 20, 158–172 (2020).

25. S. Read, V. Malmström, F. Powrie, Cytotoxic T lymphocyte-associated antigen 4 plays an essential role in the function of CD25(+)CD4(+) regulatory cells that control intestinal inflammation. J Exp Med 192, 295–302 (2000).

26. C. Asseman, S. Mauze, M. W. Leach, R. L. Coffman, F. Powrie, An essential role for interleukin 10 in the function of regulatory T cells that inhibit intestinal inflammation. J Exp Med 190, 995–1004 (1999).

27. C. I. Kingsley, M. Karim, A. R. Bushell, K. J. Wood, CD25+CD4+ regulatory T cells prevent graft rejection: CTLA-4- and IL-10-dependent immunoregulation of alloresponses. J Immunol 168, 1080–1086 (2002).

28. L. W. Collison, M. R. Pillai, V. Chaturvedi, D. A. Vignali, Regulatory T cell suppression is potentiated by target T cells in a cell contact, IL-35- and IL-10-dependent manner. J Immunol 182, 6121–6128 (2009).

29. S. A. Lim et al., Lipid signalling enforces functional specialization of Treg cells in tumors. Nature 591, 306–311 (2021).

30. S. Xu et al., Uptake of oxidized lipids by the scavenger receptor CD36 promotes lipid peroxidation and dysfunction in CD8+T cells in tumors. Immunity 54, 1561–1577.e1567 (2021).

31. H. Wang et al., CD36-mediated metabolic adaptation supports regulatory T cell survival and function in tumors. Nat Immunol 21, 298–308 (2020).

32. C. Xu et al., The glutathione peroxidase Gpx4 prevents lipid peroxidation and ferroptosis to sustain Treg cell activation and suppression of antitumor immunity. Cell Rep 35, 109235 (2021).

33. S. Tyagi, P. Gupta, A. S. Saini, C. Kaushal, S. Sharma, The peroxisome proliferator-activated receptor: A family of nuclear receptors role in various diseases. J Adv Pharm Technol Res 2, 236–240 (2011).

34. C. H. Lee, P. Olson, R. M. Evans, Minireview: lipid metabolism, metabolic diseases, and peroxisome proliferator-activated receptors. Endocrinology 144, 2201–2207 (2003).

35. R. A. Daynes, D. C. Jones, Emerging roles of PPARs in inflammation and immunity. Nat Rev Immunol 2, 748–759 (2002).

36. J. N. Feige, L. Gelman, L. Michalik, B. Desvergne, W. Wahli, From molecular action to physiological outputs: peroxisome proliferator-activated receptors are nuclear receptors at the crossroads of key cellular functions. Prog Lipid Res 45, 120–159 (2006).

37. L. Xiao, N. Wang, PPAR-δ: A key nuclear receptor in vascular function and remodeling. J Mol Cell Cardiol 169, 1–9 (2022).

38. A. K. Strosznajder, S. Wójtowicz, M. J. Jeżyna, G. Y. Sun, J. B. Strosznajder, Recent Insights on the Role of PPAR-β/δ in Neuroinflammation and Neurodegeneration, and Its Potential Target for Therapy. Neuromolecular Med 23, 86–98 (2021).

39. G. D. Barish, V. A. Narkar, R. M. Evans, PPAR delta: a dagger in the heart of the metabolic syndrome. J Clin Invest 116, 590–597 (2006).

40. S. Kanakasabai et al., Peroxisome proliferator-activated receptor delta agonists inhibit T helper type 1 (Th1) and Th17 responses in experimental allergic encephalomyelitis. Immunology 130, 572–588 (2010).

41. S. Kanakasabai, C. C. Walline, S. Chakraborty, J. J. Bright, PPARδ deficient mice develop elevated Th1/Th17 responses and prolonged experimental autoimmune encephalomyelitis. Brain Res 1376, 101–112 (2011).

42. S. E. Dunn et al., Peroxisome proliferator-activated receptor delta limits the expansion of pathogenic Th cells during central nervous system autoimmunity. J Exp Med 207, 1599–1608 (2010).

43. Y. Barak et al., Effects of peroxisome proliferator-activated receptor delta on placentation, adiposity, and colorectal cancer. Proc Natl Acad Sci U S A 99, 303–308 (2002).

44. A. E. Overacre-Delgoffe et al., Interferon-gamma drives Treg fragility to promote anti-tumor immunity. Cell 169, 1130–1141.e1111 (2017).

45. D. V. Sawant et al., Adaptive plasticity of IL-10+ and IL-35+ Treg cells cooperatively promotes tumor T cell exhaustion. Nat Immunol 20, 724–735 (2019).

46. J. L. Chao, P. A. Savage, Unlocking the Complexities of Tumor-Associated Regulatory T Cells. J Immunol 200, 415–421 (2018).

47. T. Kambayashi, T. M. Laufer, Atypical MHC class II-expressing antigen-presenting cells: can anything replace a dendritic cell? Nat Rev Immunol 14, 719–730 (2014).

48. W. Reith, S. LeibundGut-Landmann, J. M. Waldburger, Regulation of MHC class II gene expression by the class II transactivator. Nat Rev Immunol 5, 793–806 (2005).

49. D. A. Vignali, The interaction between CD4 and MHC class II molecules and its effect on T cell function. Behring Inst Mitt, 133–147 (1994).

50. E. Baixeras et al., Characterization of the lymphocyte activation gene 3-encoded protein. A new ligand for human leukocyte antigen class II antigens. J Exp Med 176, 327–337 (1992).

51. G. Pasqual et al., Monitoring T cell-dendritic cell interactions in vivo by intercellular enzymatic labelling. Nature 553, 496–500 (2018).

52. M. Schuler et al., PGC1alpha expression is controlled in skeletal muscles by PPARbeta, whose ablation results in fiber-type switching, obesity, and type 2 diabetes. Cell Metab 4, 407–414 (2006).

53. C. Baecher-Allan, E. Wolf, D. A. Hafler, MHC class II expression identifies functionally distinct human regulatory T cells. J Immunol 176, 4622–4631 (2006).

54. H. S. Ko, S. M. Fu, R. J. Winchester, D. T. Yu, H. G. Kunkel, Ia determinants on stimulated human T lymphocytes. Occurrence on mitogen- and antigen-activated T cells. J Exp Med 150, 246–255 (1979).

55. R. L. Evans et al., Peripheral human T cells sensitized in mixed leukocyte culture synthesize and express Ia-like antigens. J Exp Med 148, 1440–1445 (1978).

56. C. H. Chang, S. C. Hong, C. C. Hughes, C. A. Janeway, R. A. Flavell, CIITA activates the expression of MHC class II genes in mouse T cells. Int Immunol 7, 1515–1518 (1995).

57. J. A. León Machado, V. Steimle, The MHC Class II Transactivator CIITA: Not (Quite) the Odd-One-Out Anymore among NLR Proteins. Int J Mol Sci 22, (2021).

58. T. Wyss-Coray, C. Brander, F. Bettens, D. Mijic, W. J. Pichler, Use of antibody/peptide constructs of direct antigenic peptides to T cells: evidence for T cell processing and presentation. Cell Immunol 139, 268–273 (1992).

59. V. Barnaba, C. Watts, M. de Boer, P. Lane, A. Lanzavecchia, Professional presentation of antigen by activated human T cells. Eur J Immunol 24, 71–75 (1994).

60. M. Peiser, A. Becht, R. Wanner, Antibody blocking of MHC II on human activated regulatory T cells abrogates their suppressive potential. Allergy 62, 773–780 (2007).

61. P. Marrack, K. Rubtsova, J. Scott-Browne, J. W. Kappler, T cell receptor specificity for major histocompatibility complex proteins. Curr Opin Immunol 20, 203–207 (2008).

62. H. Yang et al., Highly immunosuppressive HLADRhi regulatory T cells are associated with unfavorable outcomes in cervical squamous cell carcinoma. Int J Cancer 146, 1993–2006 (2020).

63. R. Q. Gao et al., Circulating the HLA-DR+ T Cell Ratio Is a Prognostic Factor for Recurrence of Patients with Hepatocellular Carcinoma after Curative Surgery. J Oncol 2023, 1875153 (2023).

64. A. Machicote, S. Belén, P. Baz, L. A. Billordo, L. Fainboim, Human CD8+HLA-DR+ regulatory T cells, similarly to classical CD4+Foxp3+ cells, suppress immune responses via PD-1/PD-L1 Axis. Front Immunol 9, 2788 (2018).

65. S. Y. Park et al., Developmental defects of lymphoid cells in Jak3 kinase-deficient mice. Immunity 3, 771–782 (1995).

66. A. M. Baird, D. C. Thomis, L. J. Berg, T cell development and activation in Jak3-deficient mice. J Leukoc Biol 63, 669–677 (1998).

67. H. E. Tibbles et al., Role of a JAK3-dependent biochemical signaling pathway in platelet activation and aggregation. J Biol Chem 276, 17815–17822 (2001).

68. G. Wang et al., Regulatory effects of the JAK3/STAT1 pathway on the release of secreted phospholipase A₂-IIA in microvascular endothelial cells of the injured brain. J Neuroinflammation 9, 170 (2012).

69. D. M. Muoio et al., Fatty acid homeostasis and induction of lipid regulatory genes in skeletal muscles of peroxisome proliferator-activated receptor (PPAR) alpha knock-out mice. Evidence for compensatory regulation by PPAR delta. J Biol Chem 277, 26089–26097 (2002).

70. C. S. Loo et al., A Genome-wide CRISPR Screen Reveals a Role for the Non-canonical Nucleosome-Remodeling BAF Complex in Foxp3 Expression and Regulatory T Cell Function. Immunity 53, 143–157.e148 (2020).

71. J. van der Veeken et al., The Transcription Factor Foxp3 Shapes Regulatory T Cell Identity by Tuning the Activity of trans-Acting Intermediaries. Immunity 53, 971–984.e975 (2020).

72. P. J. Skene, S. Henikoff, An efficient targeted nuclease strategy for high-resolution mapping of DNA binding sites. Elife 6, (2017).

